# Pro- and anti-tumour activities of CD146/MCAM in breast cancer result from its heterogeneous expression and association with epithelial to mesenchymal transition

**DOI:** 10.1101/2022.12.20.521224

**Authors:** Aarren J. Mannion, Adam F. Odell, Syed Murtuza Baker, Laura C. Matthews, Pamela F. Jones, Graham P. Cook

**Author notes:** Pamela Jones and Graham Cook contributed equally to this work.

## Abstract

CD146, also known as melanoma cell adhesion molecule (MCAM), is expressed in numerous cancers and has been implicated in the regulation of metastasis. We show that CD146 negatively regulates transendothelial migration (TEM) in breast cancer. This tumour suppressor-like activity is supported by a reduction in MCAM gene expression and increased promoter methylation in tumour tissue compared to normal breast tissue. However, increased CD146/MCAM expression is associated with poor prognosis in breast cancer, a characteristic that is difficult to reconcile with inhibition of TEM by CD146 and its epigenetic silencing. Single cell transcriptome data revealed MCAM expression in multiple cell types, including the tumour vasculature and malignant epithelial cells. MCAM expressing tumour cells were in the minority and expression was associated with epithelial to mesenchymal transition (EMT). Furthermore, gene expression signatures defining invasiveness and a stem cell-like phenotype were most strongly associated with mesenchymal-like tumour cells with low levels of MCAM mRNA, likely to represent an intermediate or hybrid E/M state. Our results show that high levels of MCAM gene expression are associated with poor prognosis in breast cancer because they reflect tumour vascularisation and EMT. However, the inhibitory effects of CD146 on TEM are likely to be weakest in an intermediate state between the epithelial and mesenchymal phenotypes, consistent with highly tumourigenic nature of this population.

## Introduction

Metastatic disease is a hallmark of cancer and is responsible for the majority of cancer-related deaths (1, 2). Metastasis occurs via the infiltration of malignant cells into surrounding tissue, their entry into the lymphatic or blood vessels (intravasation) and the dissemination to distant sites, where tumour cells exit these vessels (extravasation) and seed the metastasis (3). The crossing of endothelial barriers, termed transendothelial migration (TEM), involves the interaction between the migrating cell and endothelial cells (EC) and occurs in health during inflammatory responses (4). Several studies have shown that the TEM of cancer cells occurs via very similar mechanisms to those used by extravasating leucocytes (5, 6). TEM is a multi-step process mediated by a series of receptor-ligand interactions, cytoskeletal rearrangements and migratory activity, with active participation of both the migrating cell and the endothelium. These events result in migrating cells passing between (paracellular) and through (transcellular) EC to gain access to the tissues (6, 7).

Numerous cell surface molecules expressed by both the migrating cell and EC are implicated in the regulation of TEM. For paracellular TEM, EC-EC interactions must be broken and both modes of TEM require interactions between the EC and migrating cell. The cell surface phenotype of tumour cells is thus a key factor in TEM and metastasis. Stable adhesion of cancer cells to the endothelium involves cancer cell surface molecules that are frequently over expressed in malignancy. For example, in breast cancer, MUC1 and CD44 overexpression facilitate tumour-EC interactions and promote TEM (8–11), with MUC1 overexpression linked to poor prognosis (12). Along with multicomponent EC tight junctions and adherens junctions, EC-EC contacts are regulated by CD31, CD99 and CD146 and these molecules regulate TEM of inflammatory cells (13–15). The CD146 molecule was first described as Melanoma Cell Adhesion Molecule (MCAM); this protein is highly upregulated in melanoma and was shown to mediate adhesion to EC (16). The CD146 molecule has subsequently been shown to have numerous functions in various cell types and, as such, plays a complex role in cancer progression (17).

An early step in metastasis is the generation of malignant cells with a migratory and invasive phenotype. For carcinomas, malignant epithelial cells can undergo epithelial to mesenchymal transition (EMT), differentiating them into mesenchymal-like cells which can detach from their epithelial neighbours and, having greater motility and invasive capacity, invade the surrounding tissue (18, 19). In addition, EMT promotes the acquisition of stem cell-like characteristics and drug resistance, generating cells with a potent capacity to seed metastases and resist treatment (20, 21). Not surprisingly, the expression of key genes regulating EMT are associated with patient outcomes (18, 19, 22, 23). Importantly, EMT is not simply a switch between epithelial and mesenchymal cells, but is actually a spectrum of phenotypes from fully epithelial to fully mesenchymal. Indeed, stable intermediates can be identified, known as a hybrid E/M or quasi-mesenchymal state and *in vivo* models have suggested that it is this intermediate state that contains the tumourigenic cells (21, 24–27). Furthermore, gene expression signatures characteristic of the intermediate state are markers of poor prognosis in breast cancer and several other solid tumours (28, 29).

Here we have investigated the role of CD146 in the adhesion and TEM of breast cancer cells *in vitro*. Our results suggest that CD146 expression negatively regulates these events, a conclusion supported by reduced MCAM expression in breast cancer. However, patient-based survival data suggest a pro-tumorigenic role for CD146 in breast cancer. We demonstrate that these seemingly opposing roles for CD146 can be reconciled by considering the intra-tumoural heterogeneity of breast cancer and MCAM expression and the role of EMT in generating MCAM expressing, invasive cells.

## Materials and Methods

### Cells and cell culture

Human umbilical vein ECs (HUVEC) were purchased from Promocell. Human cerebral microvascular endothelial cells (hCMEC/D3) referred to as hCMEC (VH Bio Ltd) are an immortalised cell line isolated from human temporal lobe microvessels from tissue removed to treat epilepsy (30).MDA-MB-231 cells were purchased from the European Collection of Cell Cultures and tested periodically for mycoplasma contamination. Brain metastatic derived MDA-MB-231 cells (BrM), generated by serial *in vivo* passage of MDA-MB-231 cells in mice and subsequent isolation of cancer cells from metastatic lesions of the brain (31), were a kind gift from Dr. Mihaela Lorger (University of Leeds). EC lines were cultured using endothelial cell basal medium (ECBM; Promocell), supplemented with 2% foetal calf serum (FCS) (v/v), 0.4% Endothelial Cell Growth Supplement, 0.1 ng/ml epidermal growth factor (recombinant human), 1 ng/ml basic fibroblast growth factor (recombinant human), 90 μg/ml heparin and 1 μg/ml hydrocortisone. Cells were grown on 0.2% gelatin (Sigma-Aldrich) (w/v in PBS)-coated plates. HUVEC cells were grown to passage 5 or 6. hCMEC/D3 cell lines were grown to passage 35 before discarding as cells begin to lose endothelial characteristics (30). MDA-MB-231 and BrM cells were cultured in 10% (v/v) FCS (Sigma-Aldrich)-supplemented RPMI-1640 (Sigma-Aldrich) and passaged every 3-5 days. All cell lines were incubated at 37°C under 5% CO_2_.

#### Adhesion assays

Adhesion assays were carried out as previously described (32). EC were seeded at a density of 10^4^/well of a 96-well plate (Corning) and incubated until confluent monolayers were observed. MDA-MB-231 and BrM cells were labelled with 0.4 μM Cell Tracker Green (CTG) for 30 min in serum free RPMI (SFM-RPMI) medium at 37°C. MDA-MB-231 or BrM cells were washed in SFM RPMI once before being seeded at 10^4^ per confluent EC monolayer. Adhesion assay was incubated at 37°C and CTG labelled cells were allowed to adhere to the EC monolayers for 15, 30, 60 and 120 min, after which each plate was washed once in PBS, and fixed in 4% (w/v) paraformaldehyde (PFA; Sigma-Aldrich) for 10 min, and washed twice in PBS before storage at 4°C, followed by imaging using an Incucyte Zoom Live Cell Imager (Essen Bioscience). Images were subjected to ImageJ analysis, and the ‘watershed’ function (www.imagej.net/Classic_Watershed) was used to distinguish between individual cells and clusters.

#### Transendothelial migration assay

24 well Thincert 3.0μm or 5.0μm pore diameter, transparent transwell filters (Greiner Bio-One Ltd) were coated with 0.2% (w/v) gelatin and HUVEC or hCMEC/D3 were seeded at a density of 2 ×10^4^ cells per insert. Cells were seeded in 300μl ECBM media, with 500μl in the lower chamber of the transwell insert. Endothelial cells were grown 24-48 hours to allow formation of confluent monolayers before 2 ×10^4^ breast cancer cells were seeded to the upper chamber. MDA-MB-231 and BrM cells (2 ×10^5^/ml) were CTG labelled as described in (32), before seeding to confluent EC monolayers in 1:1 ECMB:RPMI media. MDA-MB-231 and BrM cell migration was halted at 18 h by fixing in 4% PFA for 10 min followed by washing twice in 1x PBS (250μl for upper chamber and 500μl for lower chamber). Upper chambers of transwells were then scraped using cotton wool buds to remove cells on the upper layer of the transwell insert, leaving cells that had migrated to the underside of the membrane intact. Transwells were then washed twice in PBS. Migrated cells were then imaged using the EVOS microscope (Thermo Scientific).

#### Live cell imaging of intercalation

Cancer cell spreading and intercalation into endothelial monolayers is indicative of cancer cell transmigration (32–34). Intercalation was determined by live cell imaging as previously described (32, 35). Briefly, endothelial cells were seeded to 96 well plates at a density of 1×10^4^/well in 100μl to achieve confluent monolayers in 24-48 h. Once confluent endothelial monolayers were established, CTG labelled cancer cells (as described in adhesion assay) were seeded onto endothelial monolayers at a density of 1×10^4^ per well in 50μl of media (total media volume 150μl including endothelial culture medium). Plates were then imaged immediately using Live Cell Imager - Incucyte Zoom. Images were taken every 5 min for 4 h using 20x objective. Images were analysed as previously described (32).

#### RNA interference

MDA-MB-231 and BrM were transfected with SMARTpool siRNA (Dharmacon) targeting CD146 alongside a control scrambled (scr) siRNA. Transfections were performed using Lipofectamine 200 RNAiMax (Invitrogen) transfection agent and Opti-MEM I Reduced Serum Medium, GlutaMAX Supplement (Gibco) according to manufacturer’s instructions. Cells were transfected with 30 pmol siRNA in a six-well plate (2-4×10^5^ cells/well) and scaled accordingly. Briefly, for a single well of a six-well plate, 30 pmol siRNA duplexes were made in 250 μl of OptiMEM medium and incubated at room temperature for 5 min. At the same time, 5 μl Lipofectamine was made up in 250 μl of OptiMEM and incubated at room temperature for 5 min. siRNA and Lipofectamine mix were combined within the six-well plate and gently mixed before incubating at room temperature for 20 min. Following this, OptiMEM suspended cells were added to siRNA Lipofectamine complexes at 2-4×10^5^ cells in 1 ml of OptiMEM. Cells were incubated in this mixture for 4-6 h, before transfection medium was aspirated and replaced with supplemented normal culture medium. siRNA-treated cells were incubated for 24-72 h before being used in downstream assays. The siRNA molecules used (from Dharmacon/Horizon Discovery) are shown in Supplementary Table 1.

#### Flow cytometry

Cultured cells were PBS washed and trypsinised with 1× Acutase (Gibco). Cells were washed in ice-cold PBS followed by centrifugation at 300 ***g*** for 5 min. After repeated washing in PBS, cells were resuspended in 100 μl fluorescence-activated cell sorting buffer (PBS, 2% FCS and 0.09% NaN_3_) and stained with fluorophore-conjugated antibodies (CD146-APC, SHM-57 or P1H12, BioLegend), (EPCAM-FITC, B29.1, VU-ID9, Abcam), (CD44-FITC, DB105, Miltenyi Biotec), (CD99-APC, HCD99 12E7, BioLegend) and relevant isotype control antibodies at 10^6^ cells per 100 μl staining buffer for 30 min at room temperature. Stained cells were washed and fixed in Cytofix Fixation buffer (BD Biosciences) before analysis using a LSRII flow cytometer (BD Biosciences).

#### Patient sample-based gene expression, promoter methylation and survival analysis

We used two bulk tumour RNA-seq datasets in this work; one including multiple primary breast cancers, adjacent normal tissue and tissue from breast reduction surgery (36) and the other, a series of matched primary and brain metastases (37). For the latter, the normalised RNA-seq data provided by the authors was analysed directly. For the former, we downloaded metadata and raw short read archive (SRA) files (from Gene Expression Omnibus data series GSE58135), converted SRA files to FASTQ format and mapped them to human genome GRCh38 using STAR aligner v.2.5.1a. We used HTSeq v.0.10.0 to generate count matrices for genes across the samples. Raw counts were used for downstream data analysis in DESEq2; we created the DESeq2 object with raw counts with the cell metadata as the design matrix. We pre-filtered the reads that had at least 10 reads in total. To normalise, we used the median of ratio normalisation method within DESeq2 and applied variance stabilising transformation (VST) to stabilise variance across the mean. Expression data from breast cancer samples (and normal tissue) was also obtained from The Cancer Genome Atlas (TCGA) via The Cancer Immunome Atlas (38), available at https://tcia.at/home. In addition, we validated tumour versus normal tissue expression using the Gene Expression database of Normal and Tumour Tissue (GENT)2 tool (39), available at http://gent2.appex.kr/gent2/. MCAM promoter methylation was analysed using the Shiny Methylation Analysis Resource Tool (SMART) (40), available at http://www.bioinfo-zs.com/smartapp/. For the association of gene expression with patient outcomes, we used the Kaplan Meier Plotter resource which incorporates breast cancer microarray data (41) and breast cancer data from the pan-cancer RNA-seq data collection, both available at www.kmplot.com. Gene expression was compared between patient groups using the statistical tests described in the figure legends, performed using GraphPad Prism software. For single cell (sc) analysis, we utilised scRNA-seq data from a study of 26 primary breast cancers (42); data was visualised, analysed and downloaded from this study using Single Cell Portal (from the Broad Institute; https://singlecell.broadinstitute.org/single_cell). Single cell data was used to analyse expression of individual genes or to derive scores based on gene expression signatures. To sub-divide populations into MCAM^high^, MCAM^low^ or MCAM^neg^ cells, we ranked MCAM expression within a group, marked cells without MCAM gene expression and divided the remaining MCAM expressing population into two equal size groups (for odd numbers of cells, we included an additional cell in the MCAM^low^ group).

#### Gene expression signatures

To determine the relative location of individual cells or tumours on the EMT spectrum we derived an EMT score (sEMT), calculated as the sum of expression of epithelial marker genes (CDH1, GRHL2, ITGB4, KRT5, KRT8, FST) subtracted from the sum of expression of mesenchymal marker genes (CDH2, ZEB1, VIM, MMP1, FN1, TGFB1I1). The utility of this method to derive sEMT was validated by analysis of other epithelial and mesenchymal genes as described in Results. For the invasion score (sInv), we used the signature developed by Patsialou et al (43), with the exception that we only used genes shown to be upregulated in the invasive process. For the cancer stem cell score (sCSC), we used a twenty gene signature reported by Pece et al (44). Both sInv and sCSC were calculated as the mean expression of the genes in the respective signatures. An angiogenesis score (sAng), used by McDermott et al (45), was used to estimate tumour vasculature. A list of genes used to derive sEMT, sInv, sCSC and sAng are provided in Supplementary Table 2; the MMP1 gene appears in both sEMT and sCSC signatures, but otherwise the signatures are non-overlapping.

#### Statistical analysis

Statistical testing was performed using GraphPad Prism software and details of the parametric and non-parametric statistical tests used for different datasets are indicated in the figure legends. Data generated using GENT2, SMART and kmplot was analysed using their own inbuilt statisctical analysis tools (39–41).

## Results

We used a brain metastatic derivative of the triple negative breast cancer (TNBC) cell line MDA-MB-231 previously isolated from xenografts (31). This brain metastasis variant (here termed MDA-BrM) was compared to the parental cell line (termed MDA) for its ability to adhere to human endothelial cell (EC) layers. Two types of EC were tested, one comprising umbilical vein endothelial cells (HUVEC) and a second using an immortalised EC line representing human cerebral microvascular endothelial cells (hCMEC/D3), the latter acting as a model of the endothelial component of the blood brain barrier (30). The MDA and MDA-BrM cells were labelled with Cell Tracker Green (CTG) and seeded onto confluent HUVEC or hCMEC/D3 monolayers (Figure 1A) and the bound tumour cells quantified at various time points over a two-hour period (Figure 1B and C). Adhesion of both the MDA and MDA-BrM cells was greater on the HUVEC monolayers compared to the hCMEC/D3 monolayers. Overall, the adhesion of MDA and MDA-BrM to either HUVEC or hCMEC/D3 was similar, although MDA showed significantly greater adhesion to HUVECs at the 1 hr timepoint (P<0.05), whereas MDA-BrM showed significantly greater adhesion to the hCMEC/D3 cells at 1 hr (P<0.005), reflecting the tropism of these tumour cells for particular sites *in vivo*. However, these preferences were not evident after 2 hrs (Figure 1B and C).

**Figure 1:**
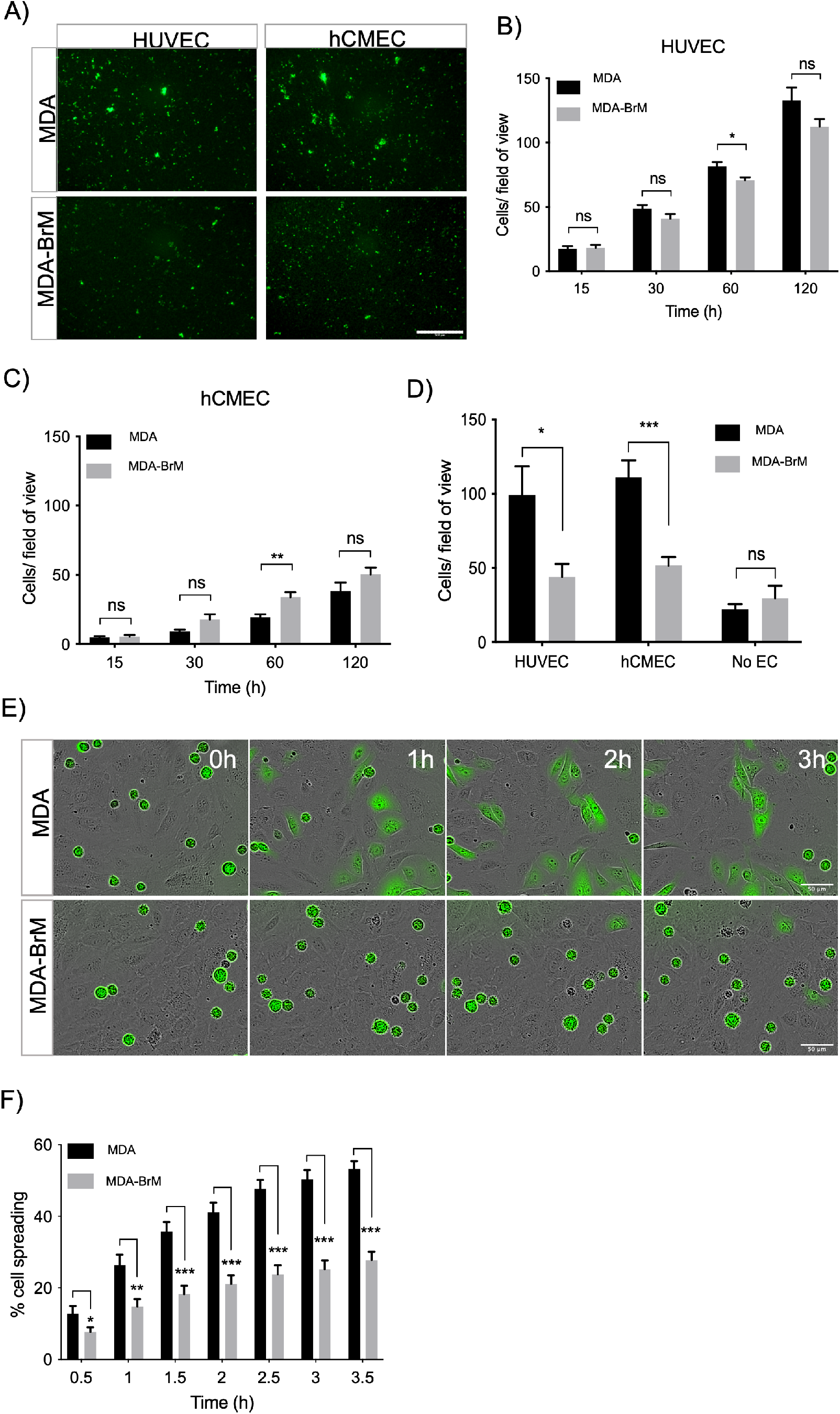
Adhesion of breast cancer cells to endothelial cells. **A)** Adhesion of CTG labelled breast cancer cell lines MDA-MB-231 (MDA) and brain metastatic variant (MDA-BrM) to HUVEC and hCMEC endothelial monolayers. Images show adhesion at 60 min time point. Scale bar: 500μm. **B)** Quantification of adhesion of MDA and MDA-BrM to HUVEC monolayers for indicated time points. Data are representative of n=3 independent experiments. Error bars indicate S.E.M. **C)** Quantification of adhesion of MDA and MDA-BrM to hCMEC monolayers for indicated time points. Data are representative of n=3 independent experiments. Error bars indicate S.E.M. **D)** Quantification of TEM of CTG labelled MDA or MDA-BrM seeded to HUVEC and hCMEC monolayers grown in the upper chamber of a Boyden transwell insert, or an empty transwell insert for No EC condition. CTG cells that had migrated to the underside of the transwell filter were imaged and quantified 18h post-seeding. Data are representative of n=3 independent experiments. Error bars indicate S.E.M. **E)** MDA and MDA-BrM TEM and intercalation determined by live cell imaging. MDA and MDA-BrM were CTG labelled and seeded to HUVEC monolayers and intercalation/spreading was captured using live cell imaging. Images were taken every 5 min for 3.5 h using a 20x objective. Scale bar: 50μm. **F)** Quantification of data in panel E, indicating the percentage of MDA or MDA-BrM cells that have undergone spreading/intercalation as a percentage of total cells. Data are representative of n=3 independent experiments. Error bars indicate S.E.M. (\**P*<0.05; \*\**P*<0.005; \*\*\**P*<0.0005; ns, not significant).

We determined whether differential adhesion of MDA and MDA-BrM to EC monolayers impacted upon the ability of these tumour cells to undergo TEM using a transwell assay; HUVEC or hCMEC/D3 cells were grown to confluency on the upper membrane of the transwell chamber and CTG-labelled MDA or MDA-BrM added in serum free media. Lower chambers of the transwell contained 10% serum, providing a migratory stimulus to the tumour cells. Following an 18 hr incubation, quantification of CTG-labelled tumour cells in the lower chamber revealed that MDA-BrM possessed significantly reduced TEM compared to the parental MDA line using both HUVEC (P<0.05) and hCMEC/D3 (P<0.0005) endothelial barriers (Figure 1D). Importantly, no significant differences in serum-stimulated migration between MDA and MDA-BrM were identified in the absence of an EC barrier, indicating that differential TEM activity of MDA and MDA-BrM was due to interactions with the endothelial cell barrier rather than intrinsic differences in migratory activity (Figure 1D).

After initial adhesion, cells undergoing TEM exhibit a morphological change and spread over the endothelium. This is followed by migration between endothelial cells, a process termed intercalation; this can be followed *in vitro* using EC monolayers and live cell imaging (32, 33). CTG-labelled MDA and MDA-BrM cells were seeded at equal density onto confluent HUVEC monolayers and imaged over a period of 4 hours to capture intercalation activity. Visual inspection of the images suggested that MDA-BrM was inferior at intercalation into HUVEC monolayers (Figure 1E) and quantification confirmed that MDA-BrM cells were significantly impaired in intercalating activity in comparison to MDA at all time points analysed (P<0.05-P<0.0005; Figure 1F). These results support the transwell migration assay data (Figure 1D), revealing that MDA-BrM has a greatly reduced capacity to undergo TEM in comparison to its parental counterpart.

A number of cell surface molecules have been implicated in the regulation of TEM, including CD44, CD99, CD155 and CD146 (10, 11, 14, 15, 47). We analysed the cell surface expression of these molecules, along with Ep-CAM, a marker of the metastatic phenotype and poor prognosis in breast cancer (48). Cell surface expression of CD44, CD99 and Ep-CAM was similar on MDA and MDA-BrM, but CD146 was expressed 8-10 fold higher on the MDA-BrM cells (P<0.005; Figure 2A and B). This difference in expression was also seen at the total protein level (Figure 2C).

**Figure 2:**
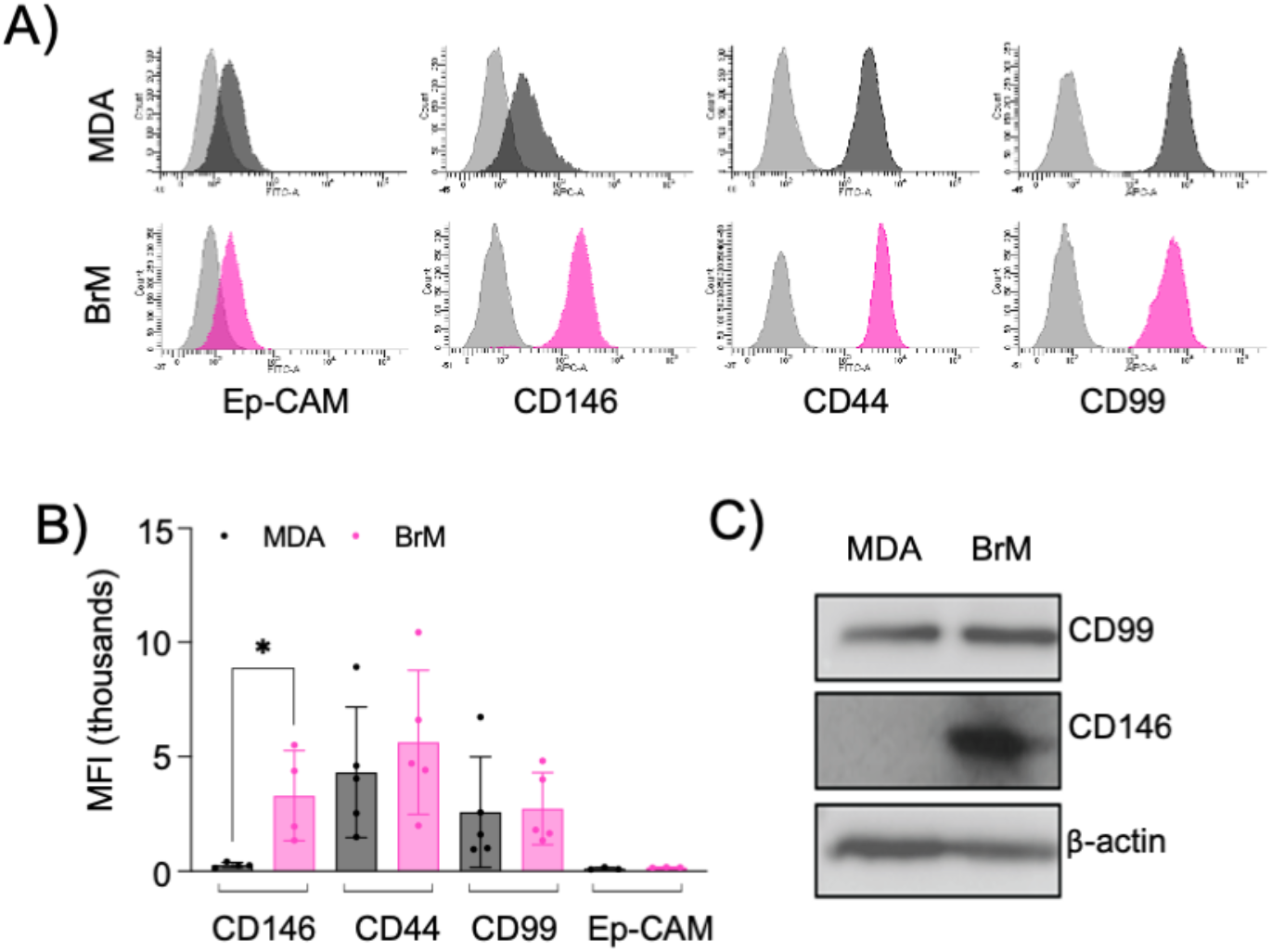
MCAM is highly expressed by brain metastatic variant of breast cancer. **A)** Expression of cell surface CD146, CD44, CD99 and Ep-CAM on MDA (dark grey histograms) and MDA-BrM (pink histograms) determined by flow cytometry compared to the isotype control (light grey histograms), representative data from *n*=5 independent experiments. **B)** Quantification of flow cytometry data shown in panel A. The graph shows quantification of indicated receptor normalised to isotype controls. *n*=5 independent experiments. Error bars indicate S.D. (\**P*<0.05). **C)** Expression of total CD146 and CD99 protein expression determined by western blotting in MDA and MDA-BrM using anti-CD146 and anti-CD99 antibody and anti-β actin as a loading control.

Cell surface CD146 participates in the TEM of inflammatory cells and melanoma cells and we speculated that it might also regulate the TEM of breast cancer cells. We used siRNA to inhibit CD146 expression and obtained a 75% reduction in cell surface CD146 in MDA-BrM cells (Figure 3A; P<0.05). The MDA cells express a ~10 fold lower level of cell surface CD146 than MDA-BrM and siRNA targeting reduced this expression by ~50% (Figure 3B; P<0.05). We labelled the siRNA transfected MDA and MDA-BrM cells with CTG and performed a HUVEC adhesion assay; for both MDA and MDA-BrM, reduced cell surface expression of CD146 was associated with significantly increased adhesion to HUVEC monolayers at the later time points in the assay (P<0.05-P<0.0005; Figure 3C and D) revealing that CD146 expression inhibits breast cancer-EC adhesion. However, the intercalation of MDA and MDA-BrM into HUVEC monolayers was unaffected by siRNA mediated changes in CD146 expression (Figure 3E). These results reveal that CD146 is important in the initial tumour-EC adhesion events but does not participate in the subsequent intercalation into the endothelial barrier.

**Figure 3:**
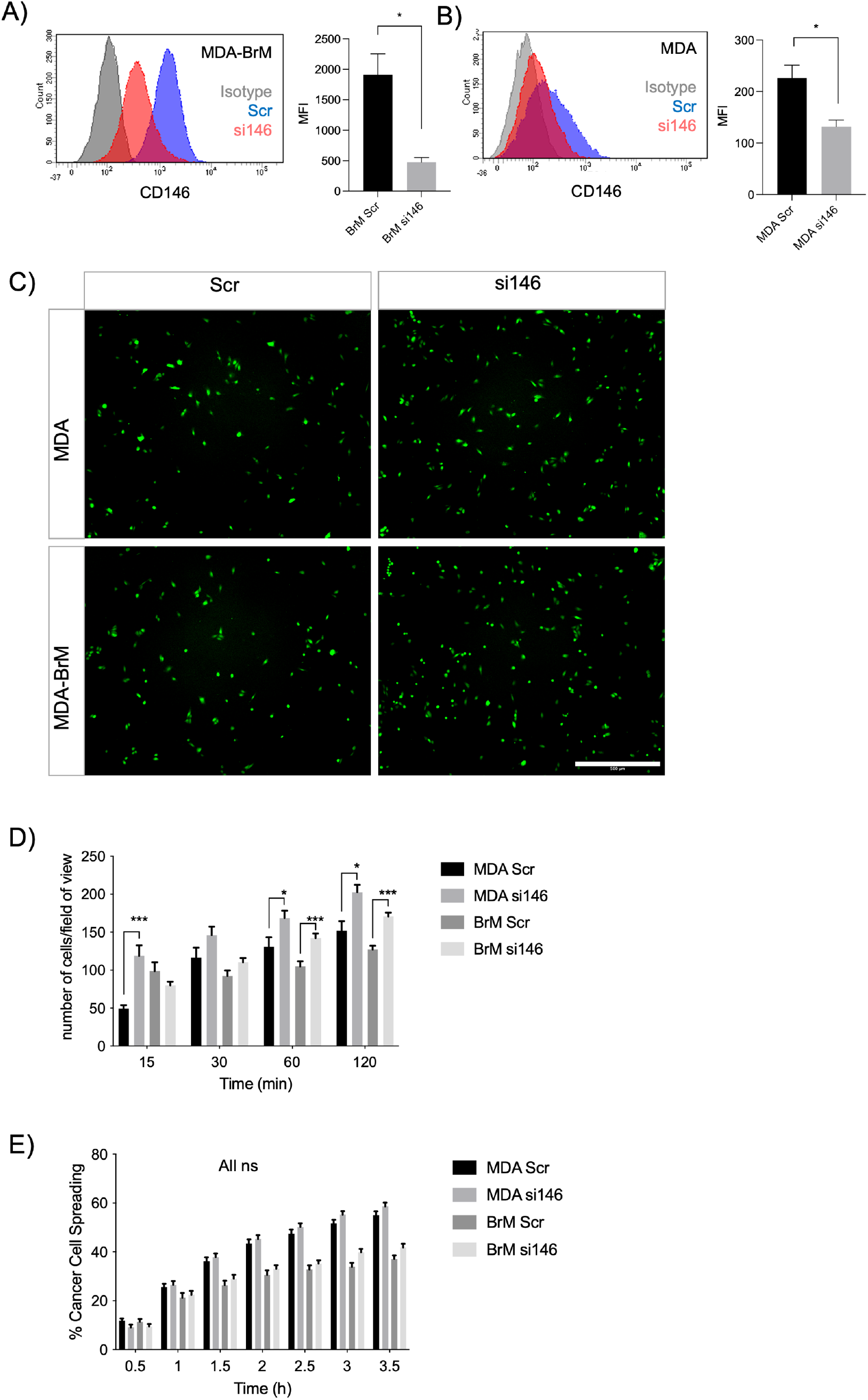
MCAM negatively regulates adhesion of breast cancer cells to endothelial cells. **A)** MDA-BrM cells were transiently transfected with siRNA targeting CD146 (si146) or a scrambled control siRNA (Scr), and CD146 expression was determined using flow cytometry 72-96 h post-transfection, compared to isotype control (grey histogram). For quantification of the flow cytometry, MFI of CD146 expression in the Scr-treated cells was compared to si146-treated cells. Data are representative of *n*=3 independent experiments. Error bars indicate S.D. (\**P*<0.05). **B)** As in panel, with MDA cells. Error bars indicate S.D. (\**P*<0.05). **C)** Images of CTG labelled MDA or MDA-BrM transfected with siRNA targeting CD146 (si146) or a scrambled control siRNA (Scr) adhering to HUVEC endothelial monolayers. Images show adhesion at 60 min time point. Scale bar: 500μm. **D)** Quantification of adhesion shown in panel C of siRNA transfected MDA or MDA-BrM with CD146 (si146) or scrambled (Scr) control siRNA MDA HUVEC monolayers for indicated time points. Data are representative of n=3 independent experiments. Error bars indicate S.E.M. **E)** Quantification of MDA and MDA-BrM transfected with CD146 (si146) or scrambled (Scr) control siRNA and subsequent TEM and intercalation determined by live cell imaging. siRNA treated MDA and MDA-BrM were CTG labelled and seeded to HUVEC monolayers and intercalation/spreading was captured using live cell imaging. Quantification indicates the percentage of MDA or MDA-BrM cells that have undergone spreading/intercalation as a percentage of total cells. Images were taken every 5 min for 3.5 h using a 20x objective. Error bars indicate S.E.M. (ns, not significant).

We performed TEM assays in Boyden chambers using these siRNA treated cells; reduced expression of CD146 did not significantly affect TEM of MDA cells in this assay (Figure 4A). However, for MDA-BrM, where unmanipulated CD146 expression was ~10 fold higher than in MDA, the reduction in CD146 expression resulted in a significant increase in TEM activity (Figure 4B; P<0.01), a phenotype readily observed from the stained cell images. However, when these Boyden chamber experiments were performed in the absence of EC, greater migratory activity was observed when CD146 expression was inhibited for both MDA and MDA-BrM, suggesting that the enhanced TEM of MDA-BrM resulting from CD146 knockdown was due to increased migratory activity rather than TEM itself (Figure 4A, B). These results show that CD146 expression inhibits the migration and TEM activity of MDA-BrM and suggests that CD146 expression functions as an inhibitor of the metastatic process in breast cancer. Indeed, reduced CD146 expression allowed MDA-BrM cells to undergo TEM at a level similar to that seen in the parental MDA line, suggesting that the low levels of CD146 expressed by MDA cells are below the threshold of inhibition of TEM, whereas the high levels of CD146 on MDA-BrM are inhibitory.

**Figure 4:**
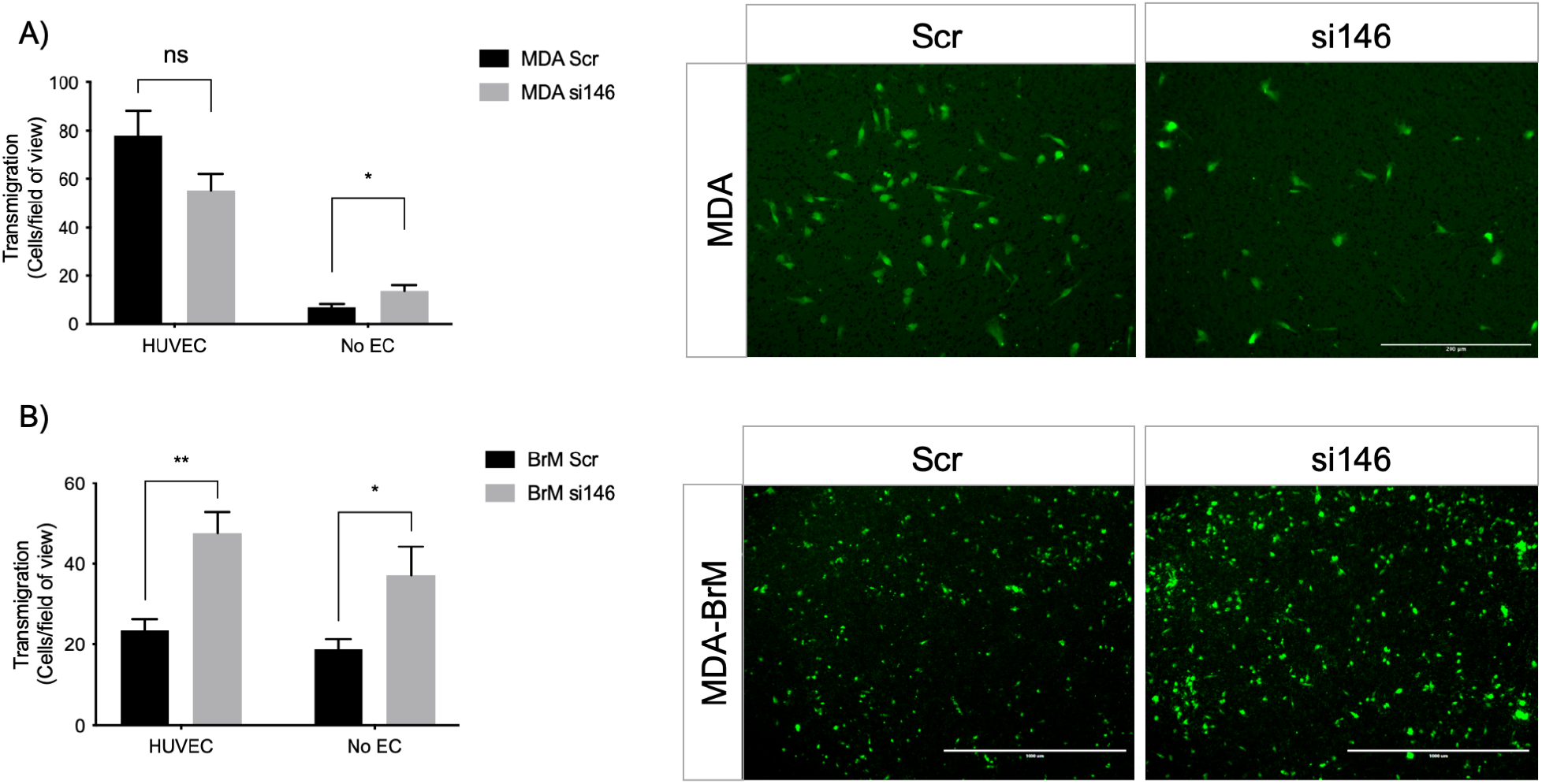
MCAM negatively regulates TEM of brain metastatic variant of breast cancer. **A)** MDA cells were transiently transfected with siRNA targeting CD146 (si146) or a scrambled control siRNA (Scr), CTG labelled, and seeded to the upper chamber of a Boyden transwell chamber in the presence of HUVEC monolayers. For the No EC condition, MDA cells were seeded to empty transwells only. CTG cells that had migrated to the underside of the transwell filter were imaged (as shown) and quantified 18h post-seeding, as shown in the bar diagrams. Scale bar: 200μm. Data are representative of n=3 independent experiments. Error bars indicate S.E.M. (\**P*<0.05; ns, not significant). **B)** As in panel A but using MDA-BrM cells. MDA-BrM were transiently transfected with siRNA targeting CD146 (si146) or a scrambled control siRNA (Scr), CTG labelled, and seeded to the upper chamber of a Boyden transwell chamber in the presence of HUVEC monolayers. For the No EC condition, MDA cells were seeded to empty transwells only. CTG cells that had migrated to the underside of the transwell filter were imaged (as shown) and quantified 18h post-seeding, as shown in the bar diagrams. Scale bar: 1000μm. Data are representative of n=3 independent experiments. Error bars indicate S.E.M. (\**P*<0.05; \*\**P*<0.005).

To address the role of CD146 expression in breast cancer progression we analysed bulk tumour transcriptome data from patient samples. We analysed the expression of the MCAM gene (encoding CD146) across a panel comprising 42 oestrogen receptor (ER)+ primary breast cancer samples, 42 primary TNBC samples and 56 samples from normal adjacent tissue or non-cancerous breast tissue remove during breast reduction surgery (36). We first characterised the samples for expression of genes which define particular breast cancer types. By definition, TNBC lack expression of ER, the progesterone receptor (PR) and HER2; we analysed expression of the cognate genes (ESR1, PRG and ERBB2 respectively) in this dataset and found that all three genes were differentially expressed in the samples as expected. Furthermore, expression of the EPCAM gene, which is overexpressed in breast cancer compared to normal tissue (48), was also differentially expressed (Supplementary Figure 1A). For the MCAM gene, we found differential expression across the three sample types (Figure 5A), with pairwise comparisons showing that MCAM expression was downregulated in both the ER+ (P<0.0001) and TNBC samples (P<0.0001) compared to the adjacent/normal breast tissue (Figure 5A). A significant reduction in MCAM gene expression in breast tumour compared to normal tissue was confirmed using two datasets from GENT2 (39), a compendium of microarray data processed to allow comparisons between studies (Supplementary Figure 1B). In addition, we analysed RNA-seq data from matched pairs of primary breast cancer and their corresponding brain metastases (37). Breast cancer brain metastases upregulate KRT13 (49) and downregulate CCDC8 (50), and these genes were differentially expressed in the primary and metastatic samples. However, MCAM expression was not significantly different between the primary tumour and the corresponding metastasis (Figure 5B).

**Figure 5:**
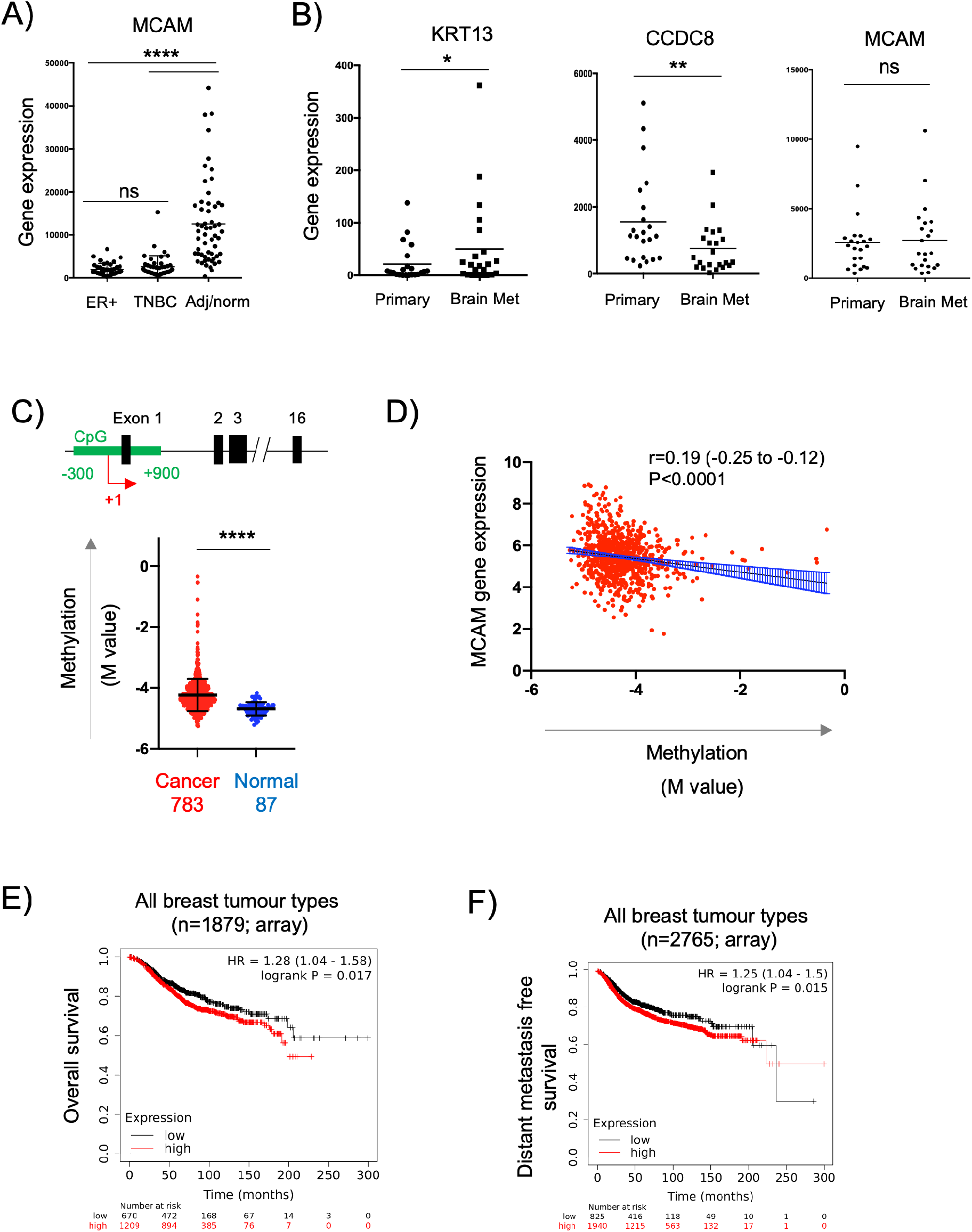
Expression of the MCAM gene in breast cancer. **A)** MCAM gene expression in ER+ breast tumours (n=42), TNBC tumours (n=42), as well as adjacent/normal breast tissue (n=56). Expression values were obtained from the data of Varley et al (36). **B)** Expression of KRT13, CCDC8 and MCAM genes in 26 primary breast cancers and patient matched brain metastases; expression values were obtained from the data of Varešlija et al. (37). **C)** Methylation of a CpG island around the transcriptional start site (TSS; +1) of the MCAM gene. The diagram shows a representation of the MCAM gene, indicating the position of the CpG island (in green) relative to the exons (black) and the TSS (+1, red arrow). The graph shows the methylation (M value) in breast cancer and normal tissue samples (793 and 87 samples respectively, from TCGA) using data from nine probes across the CpG island. For each sample, we calculated the mean M value and compared the tumour and normal tissue using a two tailed Mann Whitney test. Data analysis and download was performed using SMART (40). Methylation data for individual probes within the CpG island are shown in Supplementary Figure 2. **D)** Correlation of MCAM methylation with MCAM gene expression (using TCGA data analysed via SMART). Spearman’s r, 95% confidence internals (and associated p value) are shown together with the linear regression line and confidence intervals. **E)** Kaplan-Meier analysis of overall survival in a cohort of 1879 breast cancer patients stratified for MCAM gene expression. **F)** Kaplan-Meier analysis of distant metastasis free survival in a cohort of 2765 breast cancer patients stratified for MCAM gene expression. For E) and F), data was graphed and analysed and using KM plotter (41).

Studies using breast cancer cell lines (including MDA-MB-231) have demonstrated that the MCAM gene is regulated by promoter methylation and that treatment with demethylating agents enhances MCAM gene expression and expression of CD146 (51). This suggested that the reduced expression of the MCAM gene found in patient-derived breast cancer samples might be due to increased promoter methylation. We analysed methylation across a CpG island spanning the transcriptional start site (TSS) of MCAM using the Shiny Methylation Analysis Resource Tool (SMART), which integrates methylation and expression data from The Cancer Genome Atlas (TCGA) (40). We found significantly increased methylation in this region of the MCAM gene in cancer compared to normal tissue (Figure 5C and Supplementary Figure 2). Furthermore, MCAM gene expression was significantly and inversely correlated with methylation of this CpG island (Figure 5D).

The ability of cell surface CD146 to inhibit breast cancer TEM is consistent with the reduced expression of the MCAM gene in malignant versus normal breast tissue via epigenetic silencing. These results suggest that reduced CD146 expression in breast cancer might be a marker of poor prognosis. However, de Kruijff et al reported the opposite, finding that high CD146 expression (as determined by immunohistochemistry) is associated with reduced overall survival and reduced metastasis free survival in breast cancer (52). We performed survival analysis based on MCAM gene expression and confirmed that high MCAM gene expression was associated with significantly reduced overall survival and distant metastasis free survival when combining multiple breast cancer types (Figure 5E; P<0.05). For particular breast cancer subtypes (classified by gene expression in kmplot; 41), we found that high MCAM expression significantly reduced overall survival in HER2+ (P<0.01) and TNBC (P<0.05), but not ER+ tumours (Supplementary Figure 3A) and that high MCAM gene expression was significantly associated with a poor outcome when analysed for distant metastasis free survival in HER2+ tumours (P<0.01), but not TNBC or ER+PR+ tumours (Supplementary Figure 3B). In addition, a separate dataset (using RNAseq instead of microarray data) confirmed the association of high MCAM expression with reduced overall survival (Supplementary Figure 3C). These results mirror those of the immunohistochemistry study (52) and show that high expression of MCAM is a marker of poor prognosis and is associated with metastasis in breast cancer.

Our expression data and TEM studies suggest an anti-tumour role for CD146, whereas prognostic studies indicate a pro-tumour role. This contradiction might be explained by intra-tumoural heterogeneity of CD146/MCAM expression, with different populations of CD146 expressing cells contributing differently to disease progression. We analysed MCAM gene expression at the single cell level, using sc-RNAseq data from ~100,000 cells derived from 26 breast cancer patients (42), a dataset that includes malignant cells (~24,000), as well as normal epithelium (~4000 cells), cancer associated fibroblasts (CAF), immune cells, endothelial cells (EC) and perivascular cell (PVC) types. This dataset was viewed and analysed using Single Cell Portal at the Broad Institute. Expression of MCAM was detected in multiple cell types in breast cancer, including the malignant and normal epithelial cells, as well as other cell types, with high level expression found in EC and PVC (Figure 6A and Supplementary Figure 4). High MCAM expression in EC and PVC might account for the poor prognosis of patients with high MCAM gene expression levels, reflecting greater vascularisation of certain tumours. High expression of both the EC marker KDR/VEGFR2 and the PVC marker CSPG4 showed significant association with poor overall survival (Figure 6B; P<0.05), suggesting that high levels of MCAM gene expression reflect greater vascularisation of tumours and associated poor prognosis. This was confirmed using bulk tumour samples (1093 breast cancer patients from TCGA), where MCAM gene expression was shown to be positively correlated with an angiogenesis score (Spearman’s r=0.71; 95% CI; 0.69-0.75; Figure 6C).

**Figure 6.**
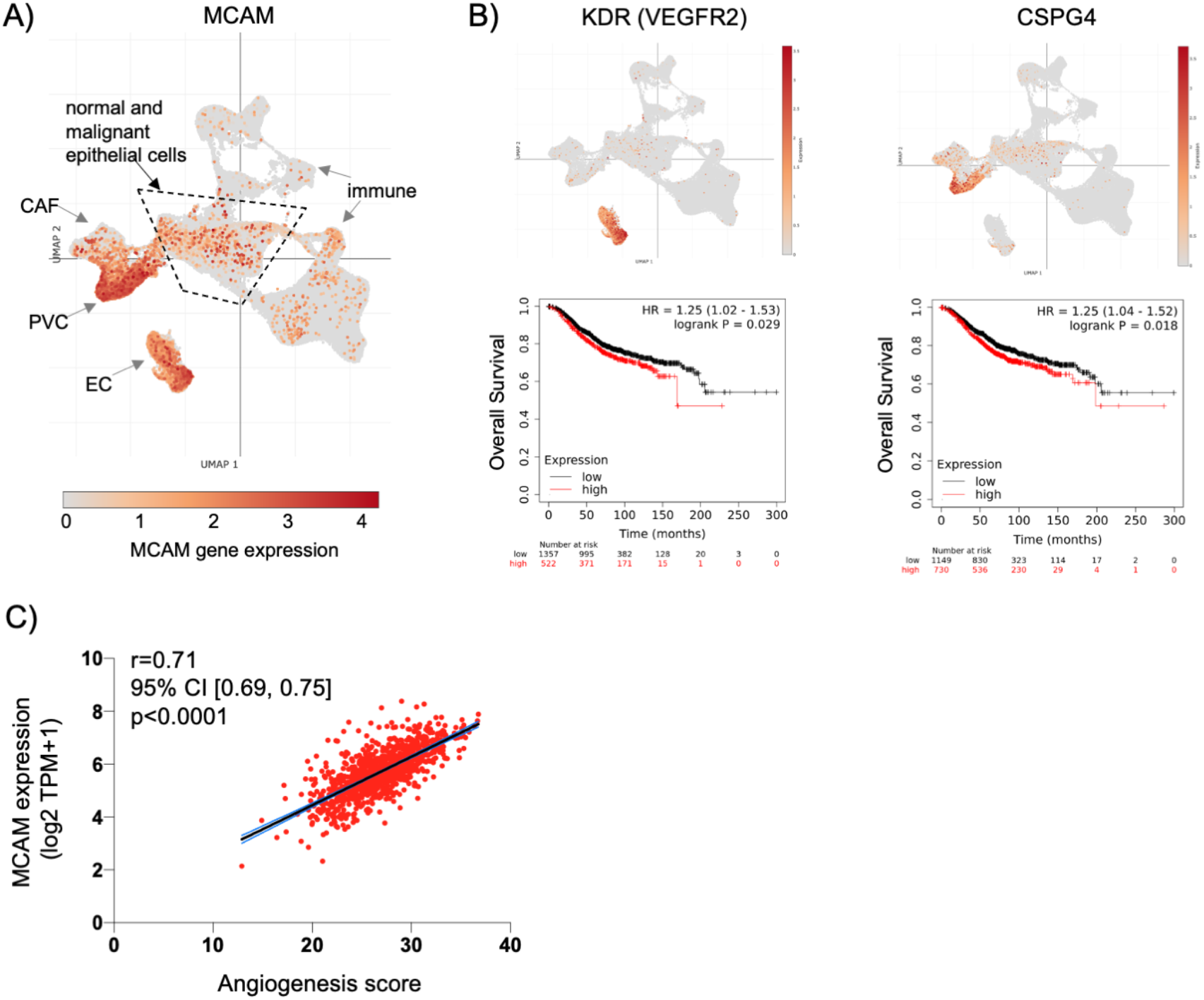
MCAM gene expression at the single cell level in human breast cancer. **A)** Uniform Manifold Approximation and Projection (UMAP) of a complete breast cancer scRNA-seq dataset comprising 100064 cells from 26 patients. This data is from the study of Wu et al with major cell populations indicated, as previously defined (42); these include immune cell types, cancer associated fibroblasts (CAF), endothelial cells (EC) and perivascular cells (PVC), as well as the epithelial cells (both malignant and normal) within the central area as indicated. Expression of MCAM is indicated and superimposed in orange. Data was displayed and analysed using Single Cell Portal. **B)** UMAP of the complete sc-RNAseq dataset showing expression of the endothelial cell marker gene KDR (VEGFR2) and the PVC marker gene CSPG4 (superimposed in orange). Below each UMAP are Kaplan Meier plots showing overall survival in 1879 breast cancer patients stratified for KDR and CSPG4 gene expression, as determined using KM plotter (41). **C)** Correlation of angiogenesis score with MCAM gene expression across a cohort of 1093 breast cancer patients from TCGA. Spearman’s r and the 95% confidence internals (with associated p value) are shown.

Our *in vitro* TEM data shows that CD146 regulates the adhesion and migration properties of the tumour cells themselves. Furthermore, high levels of tumour cell CD146 are a marker of poor outcome in breast cancer (52). We analysed MCAM gene expression within the epithelial cell populations in detail (using the sc-RNAseq data) and found that they were highly heterogenous for MCAM expression; a greater proportion of normal epithelial cells (10%) expressed MCAM transcripts at detectable levels compared to their malignant counterparts (4%). Furthermore, the expression level of MCAM was reduced in the malignant epithelial cells compared to their normal counterparts, whereas for EPCAM, the opposite relationship was found (Figure 7A). We repeated the analysis of MCAM using the sc-RNAseq data from four individual patients included in the study, choosing samples where the number of malignant cells and normal epithelial cells both exceeded one hundred. These results confirmed that MCAM expression was significantly reduced in TNBC, ER+, and ER+/HER2+ breast cancer compared to the associated normal epithelium and demonstrated intra-tumoural heterogeneity in MCAM gene expression in both normal and malignant epithelial populations (Figure 7B).

**Figure 7.**
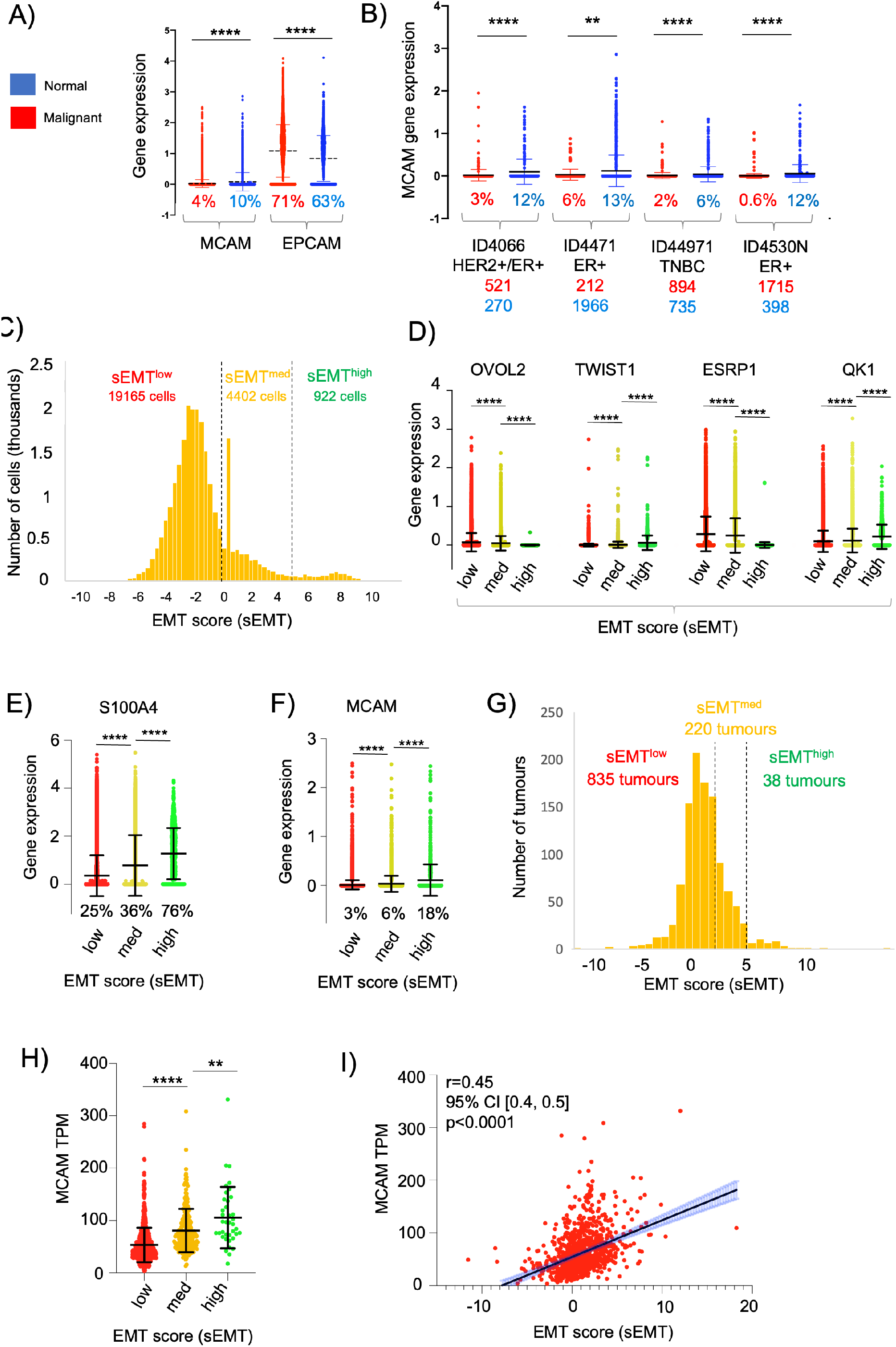
Heterogeneity of gene expression in malignant cells. **A)** MCAM and EPCAM gene expression in single malignant cells (red; n=24489) and single normal epithelial cells (blue; n=4355) as identified by Wu et al (42). Percentages indicate the proportion of expressing cells (expression>0). **B)** MCAM gene expression as in A), except performed on four individual patients from the study. Anonymised patient IDs and the phenotype of their tumour (42) are shown along with the percentage of MCAM expressing cells in the malignant (red) and normal epithelium (blue). The number of malignant and normal epithelial cells in each tumour is also shown in red and blue respectively. **C)** Population distribution of 22489 malignant cells with respect to their EMT score (sEMT). The positions of the sEMT^low^, sEMT^med^ and sEMT^high^ sub-populations are indicated, along with the number of cells in each sub-population. **D)** Expression of EMT regulators in the sEMT^low^, sEMT^med^ and sEMT^high^ sub-populations. Gene expression of the transcription factors OVOL2 and TWIST1 and the RNA splicing factors ESRP1 and QK1 are indicated. **E)** Expression of the S100A4 gene in the sEMT^low^, sEMT^med^ and sEMT^high^ populations. **F)** Expression of the MCAM gene in the sEMT^low^, sEMT^med^ and sEMT^high^ populations. **G)** Population distribution of 1093 tumours (from TCGA) with respect to their EMT score (sEMT). The positions of the sEMT^low^, sEMT^med^ and sEMT^high^ tumours are indicated, along with the numbers of tumours in each sub-group. **H)** Expression of the MCAM gene in the 1093 breast cancer tumours from TCGA defined as sEMT^low^, sEMT^med^ or sEMT^high^ tumours. **I)** Correlation of MCAM gene expression and sEMT for the in the 1093 breast cancer tumours from TCGA. Spearman’s r, confidence intervals (and associated p values) are indicated.

One important driver of intra-tumoural heterogeneity is EMT (18–21). This is a dynamic, reversible process and, within a tumour, malignant cells occupy a variety of states across the EMT spectrum rather than simply being either epithelial or mesenchymal (24, 25). We determined the relative position of each of the ~25000 malignant cells across the EMT spectrum by deriving an EMT score (sEMT) for each cell based on the expression of twelve genes, six defining the epithelial phenotype and six from the mesenchymal phenotype. This demonstrated the presence of three subpopulations based on the sEMT; sEMT^low^ (78% of cells; sEMT<0), sEMT^med^ (18%; sEMT 0-4.99) and sEMT^high^ (4%; sEMT>5), which likely represent epithelial-like cells, an intermediate population and mesenchymal-like cells respectively (Figure 7C). We analysed these three populations defined by sEMT for the expression of transcription factors which regulate EMT and for mRNA splicing factors which are differentially regulated in this differentiation pathway (19). Importantly, these genes were not used to derive the sEMT. Expression of OVOL2, which represses EMT and thus favours the epithelial phenotype, was greatest in the sEMT^low^ population and showed significantly decreasing expression in the sEMT^mid^ and sEMT^high^ cells. In contrast, expression of TWIST1, which favours the mesenchymal phenotype, increased significantly from sEMT^low^ across the three sub-populations. Similarly, expression of the epithelial splicing factor ESRP1 was significantly greater in the sEMT^low^ cells, whereas the mesenchymal splicing factor QKI was greatest in the sEMT^high^ cells (Figure 7D). The differential expression of these transcription and mRNA splicing factors validates the sEMT-based classification and supports the identification of the sEMT^low^ sub-population as epithelial-like and the sEMT^high^ population as mesenchymal-like. The intermediate levels of expression of the transcription and splicing factors in sEMT^med^ suggests that this sub-population might represent an intermediate phenotype previously termed the E/M hybrid or quasi-mesenchymal state (21, 24–27). This is further supported by the substantial and significant gain in S100A4/FSP1 expression, a marker of mesenchymal cells (53), from sEMT^med^ to sEMT^high^ (Figure 7E). We analysed MCAM expression across these sub-populations and found that MCAM expressing cells were greatly enriched in the sEMT^high^ population and expression levels increased significantly and progressively from sEMT^low^ to sEMT^med^ and sEMT^high^ (Figure 7F). This result was confirmed using bulk tumour gene expression data (1093 breast cancer patients from TCGA); there was a spectrum of sEMT across this cohort and, again, MCAM gene expression was highest in the sEMT^high^ tumours and positively correlated with sEMT (Figure 5G-I). Furthermore, both MCAM gene expression and sEMT were positively correlated with TGFB1 gene expression in the TCGA cohort (Supplementary Figure 5), consistent with the ability of TGF-β to induce EMT and MCAM gene expression (18, 54). This data suggests that heterogeneity of MCAM gene expression amongst the malignant epithelial cells in breast cancer arises, at least in part, as a result of the spectrum of EMT, both within and between tumours.

We attempted to address how levels of MCAM gene expression might be associated with the invasive and stem cell-like phenotypes that results from EMT. We derived an invasion score (sInv) and a cancer stem cell score (sCSC) for each of the ~25000 malignant cells based on published breast cancer gene expression signatures (43, 44). Not surprisingly, the sEMT^high^ population showed significantly higher sInv and sCSC than the other populations (Figure 8A and B). We sub-divided the sEMT sub-populations according to MCAM expression (high, low and no expression) and determined the mean sInv and sCSC for the nine sub-populations. Mean sInv and sCSC were strongly positively correlated, consistent with the co-acquisition of these phenotypes during EMT (Figure 8C; Spearman’s r=0.83). Importantly, the very small population of sEMT^high^MCAM^low^ cells had the greatest mean combined sInv and sCSC, suggesting that breast cancer cells of the mesenchymal-like (sEMT^high^) phenotype have the greatest invasive and stem cell potential when MCAM is expressed at low levels (Figure 8D). This is consistent with our *in vitro* data, where reduction of MCAM expression enhanced invasiveness. Separate comparisons showed that the sInv and sCSC of the sEMT^high^MCAM^low^ population were significantly greater than other sub-populations (Supplementary Figure 6A and B). Not surprisingly, the nine sub-populations showed considerable overlap with respect to sInv and sCSC, but the the sEMT^high^MCAM^low^ population had high sCSC and sInv compared to the complete population of 25,000 malignant cells (Figure 8D).

**Figure 8.**
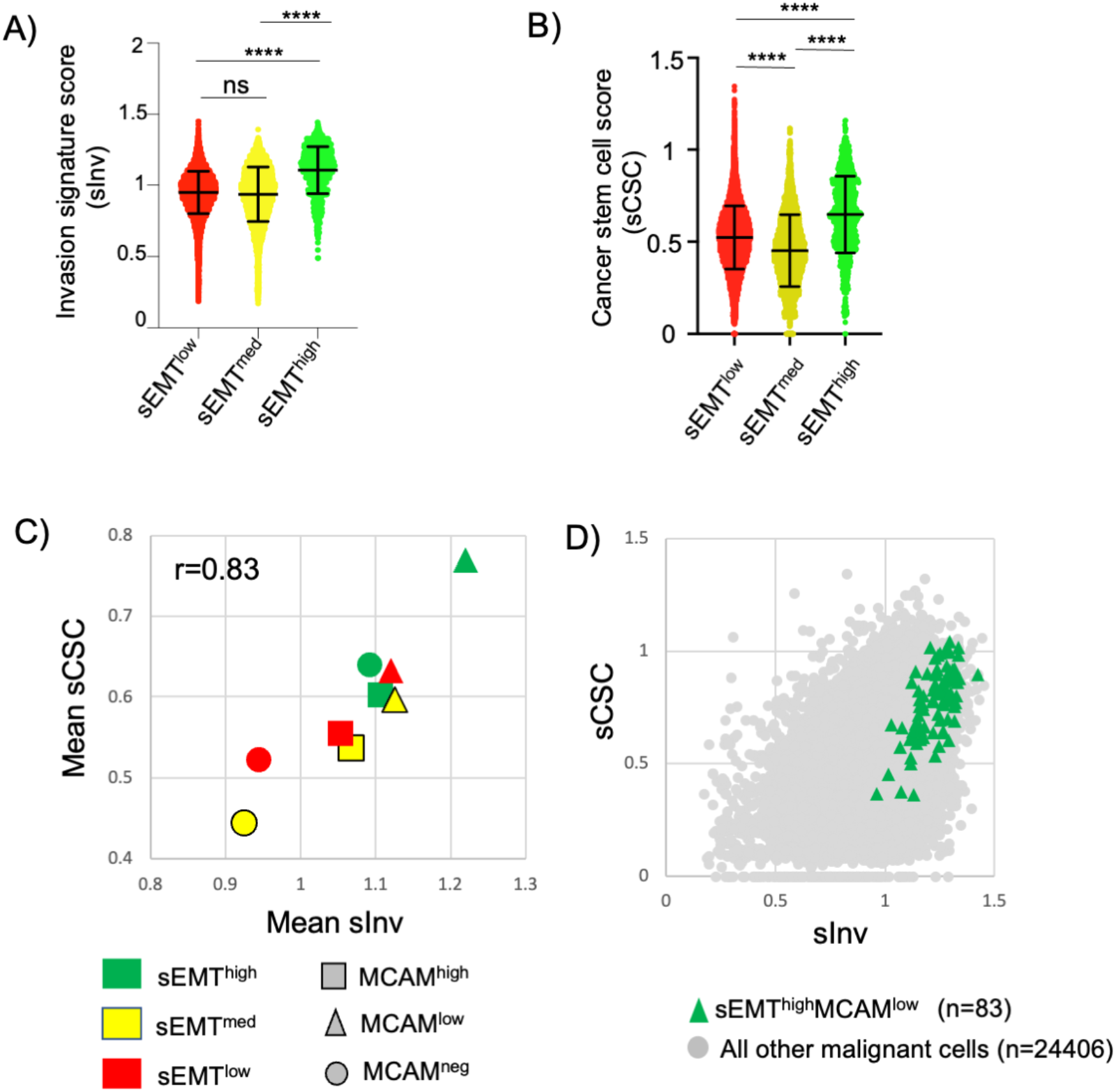
Invasive and cancer stem cell phenotypes of MCAM expressing cells. **A)** Invasion signature score (sInv) of the sEMT^low^, sEMT^med^ and sEMT^high^ sub-populations. **B)** Cancer stem cell score (sCSC) of the sEMT^low^, sEMT^med^ and sEMT^high^ sub-populations. For A) and B), the graphs show the mean and standard deviation. Statistical analysis was performed using an unpaired, two-tailed Mann Whitney test; ****p<0.0001, ns is not significant. **C)** Mean sInv and sCSC of the sEMT^low^, sEMT^med^ and sEMT^high^ sub-populations further sub-divided according to MCAM gene expression. Cells with no detectable MCAM expression were identified and then MCAM expressing cells were divided into two equal sized groups of MCAM^low^ or MCAM^high^ cells. Spearman’s r for the data is shown. **D)** Invasion signature score (sInv) and cancer stem cell score (sCSC) of the sEMT^high^MCAM^low^ sub-population (n=83) compared to all other malignant cells (n=24406).

## Discussion

Breast cancer metastasises to many sites, including bone, lung, liver and the brain and it is metastatic breast cancer that presents the major challenges to therapy (55). Our results demonstrate an inhibitory role for tumour cell-expressed CD146 in both the adhesion of breast cancer cells to EC and in migration through the endothelium. These tumour-EC interactions occur twice during metastasis, during extravasation of malignant cells from the primary tumour and again when circulating tumour cells intravasate and seed the metastasis.

Our analysis of patient transcriptome data reveals that MCAM gene expression is associated with the malignant phenotype, supporting previous findings from IHC studies of normal and malignant breast tissue (56–58). Furthermore, we show that a reduction in MCAM gene expression is associated with the increased methylation of the MCAM promoter in tumour tissue, an epigenetic modification that is known to repress gene expression and highlighted as a key regulator of breast cancer progression (59). Previous breast cancer-focussed studies have demonstrated that ectopic expression of CD146 (in MCF-7 cells) suppresses tumour growth in a xenograft model (56) and that CD146 expression is inversely correlated with Matrigel invasion (57). Collectively, these data identify CD146 as a suppressor of breast cancer progression, an activity in keeping with the epigenetic silencing of the MCAM gene in tumour tissue. Paradoxically, increased MCAM gene expression in tumour tissue was associated with reduced survival. We suggest that these seemingly contradictory findings can be reconciled by considering the heterogeneity of MCAM gene expression in breast cancer and the plasticity of the tumour phenotype.

The CD146 molecule is highly expressed in endothelium where it plays an important role in regulating extravasation. Our analysis of MCAM gene expression at the single cell level in breast cancer confirmed high levels of MCAM expression in both EC and PVC. Markers of these cell types, together with the strong positive correlation between MCAM gene expression and an angiogenesis signature, suggest that total MCAM gene expression levels in bulk tumour material are dependent, in part, on the tumour vasculature. Angiogenesis is a defining feature of solid tumours and, in common with many other cancer types, increased vascularisation is associated with poor outcomes in breast cancer (1, 60, 61).

High levels of tumour cell CD146 are also indicative of poor patient outcomes (52). Our data show that MCAM gene expression in breast cancer is indicative of the relative levels of EMT in the tumour sample. The scRNA-seq dataset used in this study is derived from 26 breast tumours (42); we classified 922 of the 24489 malignant cells as sEMT^high^. However, all but 30 of these were from a single patient (ID4513). This TNBC sample was obtained post-chemotherapy, suggesting that sEMT^high^ cells might have been enriched due to their increased drug resistance, as has been previously demonstrated (62). Although the sEMT^high^ sub-population was largely represented by this single patient in the scRNA-seq dataset, 38 of the 1093 breast tumours from the TCGA dataset were sEMT^high^ and this dataset showed a positive correlation between sEMT and MCAM gene expression. Expression of CD146 is induced by TGF-β and EMT in breast cancer cells and CD146 overexpression can drive EMT *in vitro* (54, 63, 64). When comparing EMT markers in CD146 expressing or non-expressing tumour cells, De Kruijff et al found no link between CD146 expression and EMT in breast tumours (52). However, our approach was different and we determined MCAM gene expression in cells and tumours based on their sEMT; this approach reveals that MCAM expression is highest in the more mesenchymal cells/tumours. Furthermore, our results show that heterogeneity of MCAM gene expression is found across the EMT spectrum. The epithelial phenotype predominates amongst the malignant cells and hence MCAM expressing epithelial cells are more numerous than MCAM expressing mesenchymal cells. However, the levels of MCAM expression are significantly greater in the mesenchymal-like cells, with MCAM expression levels positively correlating with sEMT. It seems likely that high levels of MCAM/CD146 expression are indicative of more cells undergoing EMT, generating larger populations of cells with invasive and stem cell-like characteristics with greater potential for metastasis and disease progression. However, further complexity in the relationship between EMT and the hallmarks of cancer is illustrated by recent findings showing that EMT is not linear, but has branchpoints with alternative outcomes (29). Interestingly, CD146 is displayed on extracellular vesicles (EV) released by the 4T1 mouse breast cancer cell line and targets the EV to the lungs where they help to establish the premetastatic niche (65), with similar EV detected in patients with breast cancer (66).

The CD146 molecule has previously been shown to play a positive role in the adhesion of melanoma cells to EC (16) and the TEM of monocytes and T cells (15, 67, 68). However, for breast cancer, our data argue for an inhibitory action of CD146 in tumour cell-EC adhesion and TEM, supporting previous work revealing that CD146 has tumour suppressor-like activity (56–58). Gene signatures revealed that mesenchymal-like cells (EMT^high^) had the highest invasive (sInv) and stemness (sCSC) scores, as expected given the well-established links between EMT, stemness and invasion. However, MCAM^low^ expressing cells had greater sInv and sCSC than MCAM^high^ expressing cells. It is tempting to speculate that intermediate levels of MCAM gene expression characterise the hybrid E/M state, a population that is greatly enriched in cells with metastatic and cancer-initiating activity (21, 24–28). Importantly, CD146 is more than a marker of EMT and overexpression can drive EMT (63). The MDA-MB-231 cell line used in our studies has a mesenchymal-like phenotype (23) and it is possible that CD146 inhibition in this cell line pushes the phenotype towards the hybrid E/M state.

The inhibitory activity of CD146 in adhesion and TEM were most pronounced in the brain metastasising variant MDA-BrM. Enhanced CD146 expression on MDA-BrM was associated with the reduced TEM phenotype using HUVEC and hCMEC/D3 cells as a source of EC from the peripheral circulation and blood brain barrier respectively. We speculated that MDA-BrM might demonstrate stronger binding to hCMEC/D3 than HUVEC and that the parental line would exhibit a preference for HUVEC, consistent with their tropism in *in vivo* models. Whilst there was some evidence of this selectivity at a single time point, this was not evident throughout the assay. Indeed, both the parental MDA and MDA-BrM derivative cell lines showed only weak binding to hCMEC/D3. This may reflect findings suggesting that adhesion to blood brain barrier EC is very weak in the absence of inflammation and that TEM at this site might be regulated differently to restrict the influx of immune cells into the brain (69). Alternatively, weak adhesion to hCMEC/D3 might reflect immortalisation by SV40/hTERT, resulting in differences between this cell line and primary blood brain barrier cells (30, 70–72).

In summary, expression of the CD146 molecule in breast cancer is of prognostic and functional importance. High levels of CD146 expression in bulk tumour reflect vascularisation and CD146 expression in the malignant cells is associated with EMT and increased invasive and stemness characteristics. Intermediate levels of MCAM gene expression are likely to be associated with the hybrid E/M state, whereas cells expressing high levels of CD146 are likely to be fully mesenchymal and have less metastatic activity *in vivo*. Our findings have relied extensively on informatics-based approaches using human breast cancer transcriptome profiles. Gene signatures underestimate the complexity of biological systems and, whilst they are valuable to infer phenotypes, they are an imperfect approach. In addition, the use of cut-offs (e.g. in gene expression or signature scores) is arbitrary with respect to biological effects and it is important to now test these hypotheses and verify key findings in biological model systems. Our results demonstrate that understanding cellular and molecular heterogeneity in breast cancer is essential to understand and treat the underlying pathology.

## Acknowledgements

We are grateful to our colleagues Mihaela Lorger and Tom Hughes for providing advice and reagents and to Tom Hughes for comments on the manuscript. We are grateful to the University of Leeds School of Medicine for scholarship support to AM.

## Author contribution statement

The study was conceived and designed by AM, AO, LCM, PJF and GPC. AM and AO performed the *in vitro* experiments, LCM, SMB and GPC performed the *in silico* analyses and all authors contributed to the drafting and editing of the manuscript. Some of the data presented in this manuscript were reported in the PhD thesis of AM, submitted to the University of Leeds.

## References

1. D. Hanahan, R. A. Weinberg, Hallmarks of cancer: The next generation. Cell 144, 646–674 (2011).

2. H. Dillekås, M. S. Rogers, O. Straume, Are 90% of deaths from cancer caused by metastases? Cancer Med. 8, 5574–5576 (2019).

3. A. W. Lambert, D. R. Pattabiraman, R. A. Weinberg, Emerging Biological Principles of Metastasis. Cell 168, 670–691 (2017).

4. D. Vestweber, How leukocytes cross the vascular endothelium. Nat. Rev. Immunol. 15, 692–704 (2015).

5. C. D. Madsen, E. Sahai, Cancer Dissemination-Lessons from Leukocytes. Dev. Cell 19, 13–26 (2010).

6. N. Reymond, B. B. D’Água, A. J. Ridley, Crossing the endothelial barrier during metastasis. Nat. Rev. Cancer 13, 858–870 (2013).

7. Z. Mamdouh, A. Mikhailov, W. A. Muller, Transcellular migration of leukocytes is mediated by the endothelial lateral border recycling compartment. J Exp Med 206, 2795–2808 (2009).

8. J. J. Rahn, et al., MUC1 mediates transendothelial migration in vitro by ligating endothelial cell ICAM-1. Clin. Exp. Metastasis 22, 475–483 (2005).

9. L. G. Yu, et al., Galectin-3 interaction with Thomsen-Friedenreich disaccharide on cancer-associated MUC1 causes increased cancer cell endothelial adhesion. J. Biol. Chem. 282, 773–781 (2007).

10. K. Zen, et al., CD44v4 is a major E-selectin ligand that mediates breast cancer cell transendothelial migration. PLoS One 3, e1826 (2008).

11. H. S. Wang, et al., CD44 Cross-linking induces integrin-mediated adhesion and transendothelial migration in breast cancer cell line by up-regulation of LFA-1 (αLβ2) and VLA-4 (α4β1). Exp. Cell Res. 304, 116–126 (2005).

12. X. Jing, H. Liang, C. Hao, X. Yang, X. Cui, Overexpression of MUC1 predicts poor prognosis in patients with breast cancer. Oncol. Rep. 41, 801–810 (2019).

13. L. Piali, et al., CD31/PECAM-1 is a ligand for αvβ3 integrin involved in adhesion of leukocytes to endothelium. J. Cell Biol. 130, 451–460 (1995).

14. A. R. Schenkel, Z. Mamdouh, X. Chen, R. M. Liebman, W. A. Muller, CD99 plays a major role in the migration of monocytes through endothelial junctions. Nat. Immunol. 3, 143–150 (2002).

15. N. Bardin, et al., CD146 and its soluble form regulate monocyte transendothelial migration. Arterioscler. Thromb. Vasc. Biol. 29, 746–753 (2009).

16. S. Xie, et al., Expression of MCAM/MUC18 by human melanoma cells leads to increased tumor growth and metastasis. Cancer Res. 57, 2295–2303 (1997).

17. Z. Wang, X. Yan, CD146, a multi-functional molecule beyond adhesion. Cancer Lett. 330, 150–162 (2013).

18. S. Lamouille, J. Xu, R. Derynck, Molecular mechanisms of epithelial-mesenchymal transition. Nat. Rev. Mol. Cell Biol. 15, 178–196 (2014).

19. A. W. Lambert, R. A. Weinberg, Linking EMT programmes to normal and neoplastic epithelial stem cells. Nat. Rev. Cancer 21, 325–338 (2021).

20. S. A. Mani, et al., The Epithelial-Mesenchymal Transition Generates Cells with Properties of Stem Cells. Cell 133, 704–715 (2008).

21. T. Celià-Terrassa, M. K. Jolly, Cancer stem cells and epithelial-to-mesenchymal transition in cancer metastasis. Cold Spring Harb. Perspect. Med. 10, 1–17 (2020).

22. S. E. Moody, et al., The transcriptional repressor Snail promotes mammary tumor recurrence. Cancer Cell 8, 197–209 (2005).

23. J. H. Taube, et al., Core epithelial-to-mesenchymal transition interactome gene-expression signature is associated with claudin-low and metaplastic breast cancer subtypes. Proc. Natl. Acad. Sci. U. S. A. 107, 15449–15454 (2010).

24. I. Pastushenko, et al., Identification of the tumour transition states occurring during EMT. Nature 556, 463–468 (2018).

25. C. Kröger, et al., Acquisition of a hybrid E/M state is essential for tumorigenicity of basal breast cancer cells. Proc. Natl. Acad. Sci. U. S. A. 116, 7353–7362 (2019).

26. F. Lüönd, et al., Distinct contributions of partial and full EMT to breast cancer malignancy. Dev. Cell 56, 3203–3221.e11 (2021).

27. I. Pastushenko, et al., Fat1 deletion promotes hybrid EMT state, tumour stemness and metastasis. Nature 589, 448–455 (2021).

28. A. P. Deshmukh, et al., Identification of EMT signaling cross-talk and gene regulatory networks by single-cell RNA sequencing. Proc. Natl. Acad. Sci. U. S. A. 118, e2102050118 (2021).

29. Y. Zhang, et al., Genome-wide CRISPR screen identifies PRC2 and KMT2D-COMPASS as regulators of distinct EMT trajectories that contribute differentially to metastasis. Nat. Cell Biol. 24, 554–564 (2022).

30. B. B. Weksler, et al., Blood-brain barrier-specific properties of a human adult brain endothelial cell line. FASEB J. 19, 1872–1874 (2005).

31. T. Yoneda, P. J. Williams, T. Hiraga, M. Niewolna, R. Nishimura, A bone-seeking clone exhibits different biological properties from the MDA-MB-231 parental human breast cancer cells and a brain-seeking clone in vivo and in vitro. J. Bone Miner. Res. 16, 1486–1495 (2001).

32. A. J. Mannion, A. F. Odell, A. Taylor, P. F. Jones, G. P. Cook, Tumour cell CD99 regulates transendothelial migration via CDC42 and actin remodelling. J. Cell Sci. 134, jcs240135 (2021).

33. N. Reymond, et al., Cdc42 promotes transendothelial migration of cancer cells through β1 integrin. J. Cell Biol. 199, 653–668 (2012).

34. M. D. Onken, J. Li, J. A. Cooper, Uveal Melanoma Cells Utilize a Novel Route for Transendothelial Migration. PLoS One 9, e115472 (2014).

35. A. J. Mannion, Live Cell Imaging and Analysis of Cancer-Cell Transmigration Through Endothelial Monolayers. Methods Mol. Biol. 2441, 329–338 (2022).

36. K. E. Varley, et al., Recurrent read-through fusion transcripts in breast cancer. Breast Cancer Res. Treat. 146, 287–297 (2014).

37. D. Varešlija, et al., Transcriptome Characterization of Matched Primary Breast and Brain Metastatic Tumors to Detect Novel Actionable Targets. J. Natl. Cancer Inst. 111, 388–398 (2019).

38. P. Charoentong, et al., Pan-cancer Immunogenomic Analyses Reveal Genotype-Immunophenotype Relationships and Predictors of Response to Checkpoint Blockade. Cell Rep. 18, 248–262 (2017).

39. S. J. Park, B. H. Yoon, S. K. Kim, S. Y. Kim, GENT2: An updated gene expression database for normal and tumor tissues. BMC Med. Genomics 12, Suppl 5 101 (2019).

40. Y. Li, D. Ge, C. Lu, The SMART App: An interactive web application for comprehensive DNA methylation analysis and visualization. Epigenetics and Chromatin 12, 71 (2019).

41. B. Györffy, et al., An online survival analysis tool to rapidly assess the effect of 22,277 genes on breast cancer prognosis using microarray data of 1,809 patients. Breast Cancer Res. Treat. 123, 725–731 (2010).

42. S. Z. Wu, et al., A single-cell and spatially resolved atlas of human breast cancers. Nat. Genet. 53, 1334–1347 (2021).

43. A. Patsialou, et al., Selective gene-expression profiling of migratory tumor cells in vivo predicts clinical outcome in breast cancer patients. Breast Cancer Res. 14, R319 (2012).

44. S. Pece, et al., Identification and clinical validation of a multigene assay that interrogates the biology of cancer stem cells and predicts metastasis in breast cancer: A retrospective consecutive study. EBioMedicine 42, 352–362 (2019).

45. D. F. McDermott, et al., Clinical activity and molecular correlates of response to atezolizumab alone or in combination with bevacizumab versus sunitinib in renal cell carcinoma. Nat. Med. 24, 749–757 (2018).

46. N. Reymond, P. Riou, A. J. Ridley, Rho GTPases and cancer cell transendothelial migration. Methods Mol. Biol. 827, 123–142 (2012).

47. N. Reymond, et al., DNAM-1 and PVR regulate monocyte migration through endothelial junctions. J Exp Med 199, 1331–1341 (2004).

48. B. T. F. van der Gun, et al., EpCAM in carcinogenesis: The good, the bad or the ugly. Carcinogenesis 31, 1913–1921 (2010).

49. Q. Li, et al., Keratin 13 expression reprograms bone and brain metastases of human prostate cancer cells. Oncotarget 7, 84645–84657 (2016).

50. R. P. Pangeni, et al., The GALNT9, BNC1 and CCDC8 genes are frequently epigenetically dysregulated in breast tumours that metastasise to the brain. Clin. Epigenetics 7, 57 (2015).

51. P. Dudzik, et al., Aberrant promoter methylation may be responsible for the control of CD146 (MCAM) gene expression during breast cancer progression. Acta Biochim. Pol. 66, 619–625 (2019).

52. I. E. De Kruijff, et al., The prevalence of CD146 expression in breast cancer subtypes and its relation to outcome. Cancers (Basel). 10, 134 (2018).

53. X. Ye, et al., Upholding a role for EMT in breast cancer metastasis. Nature 547, E1–E3 (2017).

54. Y. Ma, et al., CD146 mediates an E-cadherin-to-N-cadherin switch during TGF-β signaling-induced epithelial-mesenchymal transition. Cancer Lett. 430, 201–214 (2018).

55. B. Weigelt, J. L. Peterse, L. J. Van’t Veer, Breast cancer metastasis: Markers and models. Nat. Rev. Cancer 5, 591–602 (2005).

56. I. M. Shih, M. Y. Hsu, J. P. Palazzo, M. Herlyn, The cell-cell adhesion receptor Mel-CAM acts as a tumor suppressor in breast carcinoma. Am. J. Pathol. 151, 745–751 (1997).

57. A. Ouhtit, M. E. Abdraboh, A. D. Hollenbach, H. Zayed, M. H. G. Raj, CD146, a novel target of CD44-signaling, suppresses breast tumor cell invasion. Cell Commun. Signal. 15, 45 (2017).

58. G. Chakraborty, H. Rangaswami, S. Jain, G. C. Kundu, Hypoxia regulates cross-talk between Syk and Lck leading to breast cancer progression and angiogenesis. J. Biol. Chem. 281, 11322–11331 (2006).

59. B. Pasculli, R. Barbano, P. Parrella, Epigenetics of breast cancer: Biology and clinical implication in the era of precision medicine. Semin. Cancer Biol. 51, 22–35 (2018).

60. N. Weidner, J. P. Semple, W. R. Welch, J. Folkman, Tumor Angiogenesis and Metastasis — Correlation in Invasive Breast Carcinoma. N. Engl. J. Med. 324, 1–8 (1991).

61. E. R. Horak, et al., Angiogenesis, assessed by platelet/endothelial cell adhesion molecule antibodies, as indicator of node metastases and survival in breast cancer. Lancet 340, 1120–1124 (1992).

62. C. J. Creighton, et al., Residual breast cancers after conventional therapy display mesenchymal as well as tumor-initiating features. Proc. Natl. Acad. Sci. U. S. A. 106, 13820–13825 (2009).

63. Q. Zeng, et al., CD146, an epithelial-mesenchymal transition inducer, is associated with triple-negative breast cancer. Proc. Natl. Acad. Sci. U. S. A. 109, 1127–1132 (2012).

64. A. M. Imbert, et al., CD146 Expression in Human Breast Cancer Cell Lines Induces Phenotypic and Functional Changes Observed in Epithelial to Mesenchymal Transition. PLoS One 7, e43752 (2012).

65. S. Ghoroghi, et al., Ral GTPases promote breast cancer metastasis by controlling biogenesis and organ targeting of exosomes. Elife 10, 1–29 (2021).

66. K. Ekström, et al., Characterization of surface markers on extracellular vesicles isolated from lymphatic exudate from patients with breast cancer. BMC Cancer 22, 50 (2022).

67. J. Breuer, et al., Blockade of MCAM/CD146 impedes CNS infiltration of T cells over the choroid plexus. J. Neuroinflammation 15, 236 (2018).

68. H. Duan, et al., Targeting endothelial CD146 attenuates neuroinflammation by limiting lymphocyte extravasation to the CNS. Sci. Rep. 3, 1687 (2013).

69. B. Engelhardt, R. M. Ransohoff, Capture, crawl, cross: The T cell code to breach the blood-brain barriers. Trends Immunol. 33, 579–589 (2012).

70. B. Weksler, I. A. Romero, P. O. Couraud, The hCMEC/D3 cell line as a model of the human blood brain barrier. Fluids Barriers CNS 10, 16 (2013).

71. E. A. L. M. Biemans, L. Jäkel, R. M. W. de Waal, H. B. Kuiperij, M. M. Verbeek, Limitations of the hCMEC/D3 cell line as a model for Aβ clearance by the human blood-brain barrier. J. Neurosci. Res. 95, 1513–1522 (2017).

72. E. Urich, S. E. Lazic, J. Molnos, I. Wells, P. O. Freskgård, Transcriptional profiling of human brain endothelial cells reveals key properties crucial for predictive in vitro blood-brain barrier models. PLoS One 7, e38149 (2012).

