## Supplementary Figures and Legends for "Pro- and anti-tumour activities of CD146/MCAM in breast cancer result from its heterogeneous expression and association with epithelial to mesenchymal transition"

### 1. Supplementary Figures and Legends

#### Supplementary Figure 1

**A)** Verification of expression of ESR1 (oestrogen receptor), PGR1 (progesterone receptor), ERBB2 (HER2) and EPCAM genes in the ER+ breast cancers, TNBC and adjacent normal breast tissue from the study of Varley et al. P values indicate a one-way ANOVA test to test for significant variance between the means of the samples.

**B)** MCAM gene expression in breast cancer versus normal tissue as determined using the GENT2 database (39). Metadata from two different microarray platforms are indicated, with numbers of samples of tumour (red) and normal tissue (blue) indicated. ****P<0.0001 as determined by GENT2 using a two tailed Mann Whitney test.

#### Supplementary Figure 2.

Methylation of the MCAM promoter. A graphical representation of the MCAM gene on chromosome 11q23, highlighting exons, the transcriptional start site (TSS; red arrow) and a CpG island located from approximately -300 bp to +900 bp relative to the TSS. The precise sequence position of the CpG island on chromosome 11 is from positions 119317436 bp to 119316218 bp (shown in green), with the TSS located at 119317130 bp (red); this diagram is based on information from the hg38 assembly of the human genome available via the UCSC genome browser at https://genome-euro.ucsc.edu. Methylation data was obtained from nine probes within the CpG island, five upstream of the TSS (p9-p5) and four downstream (p4-p1) as indicated. The graphs show methylation levels (M values) in tumour and normal tissue for these nine probes, the genomic position of each probe is also indicated on the graph. The final graph shows aggregate data (the mean M value across the nine probes for tumour and normal tissue samples). The statistical analysis shown here was that obtained using the default setting from the SMART tool (40).

**Supplementary Figure 3.**

Survival data in breast cancer based on MCAM gene expression. A) Overall survival and B) distant metastasis free survival in HER2+ tumours, TNBC and ER+PR+ tumours stratified for MCAM gene expression. Data in A) and B) is based on microarray platforms. C) Overall survival in all breast cancer types according to MCAM gene expression in 2976 samples as determined by RNA-seq. Data was analysed and displayed using KM plotter (41).

**Supplementary Figure 4.**

MCAM gene expression in the major cell populations identified in the 26 patient/100064 cell scRNA-seq study of Wu et al (42). This data was obtained and graphed using Single Cell Portal.

**Supplementary Figure 5.**

Correlations between **A)** sEMT and TGFB1 gene expression and **B)** MCAM and TGFB1 gene expression in 1093 breast cancer patients from TCGA. Spearman's r and 95% confidence intervals (with associated p values) are shown.

**Supplementary Figure 6.**

**A)** Invasion score (sInv) versus cancer stem cell score (sCSC) for $sEMT^{low}$, $sEMT^{med}$ and $sEMT^{high}$ malignant cells, with position of $MCAM^{neg}$ (grey), $MCAM^{low}$ (green) and $MCAM^{high}$ (red) cells identified.

**B)** Invasion score (sInv) of $sEMT^{low}$, $sEMT^{med}$ and $sEMT^{high}$ populations subdivided into $MCAM^{neg}$, $MCAM^{low}$ and $MCAM^{high}$ cells. Subsets are ordered on the vertical axis based on their mean sInv. A Mann Whitney test indicated that the sEMT$^{high}$MCAM$^{low}$ population has a significantly greater sInv than any other subset.

**C)** As in B), except the subsets were scored for sCSC.

**Supplementary Figure 1**

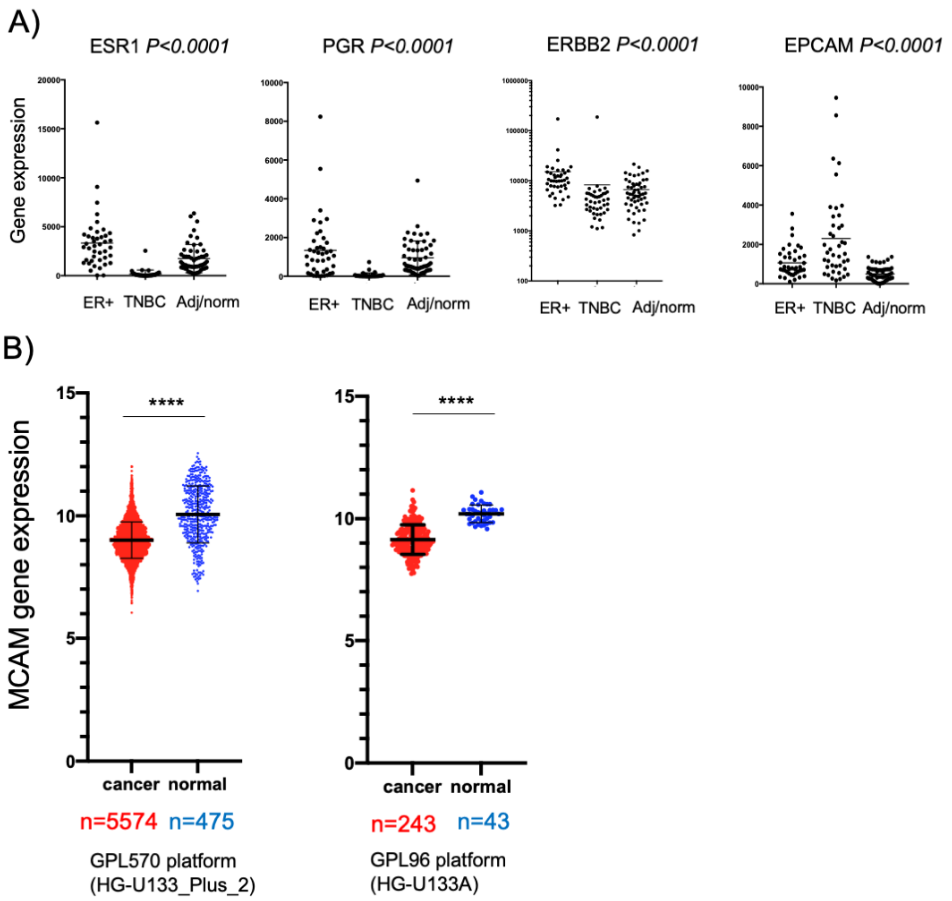

### Supplementary Figure 2

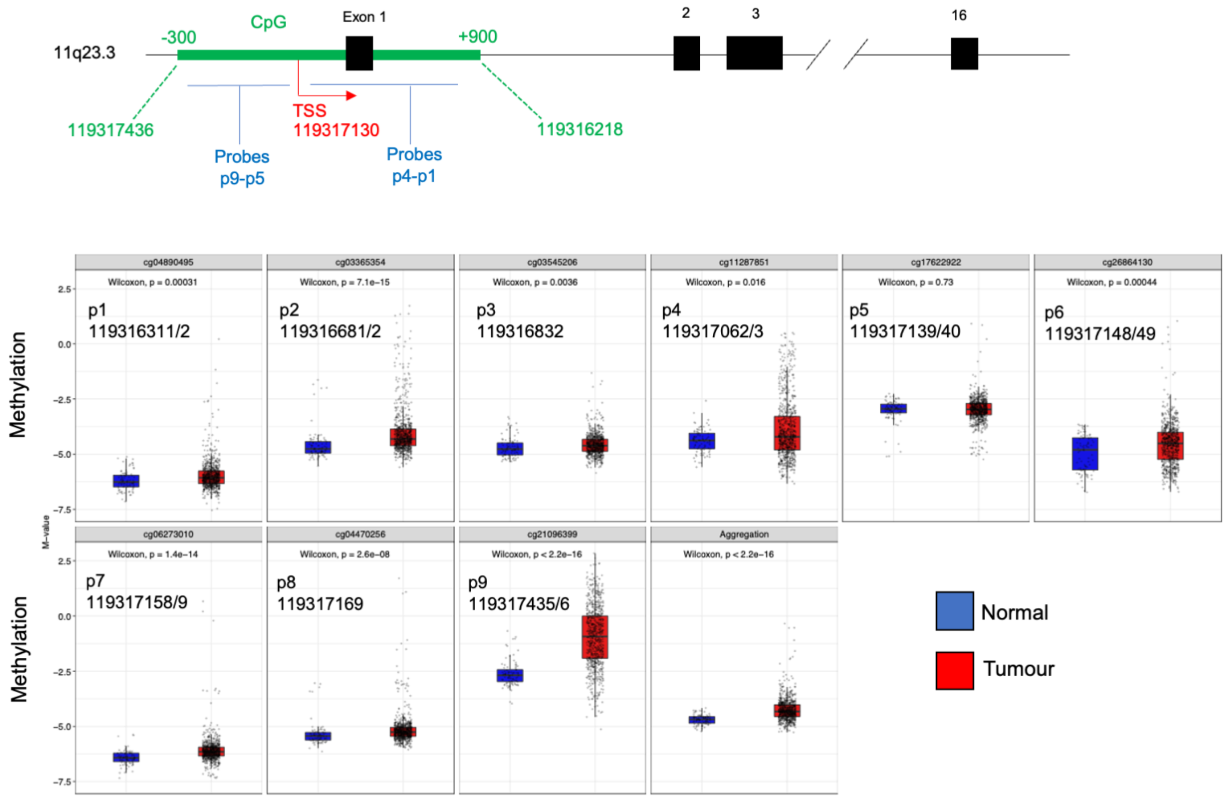


### Supplementary Figure 3

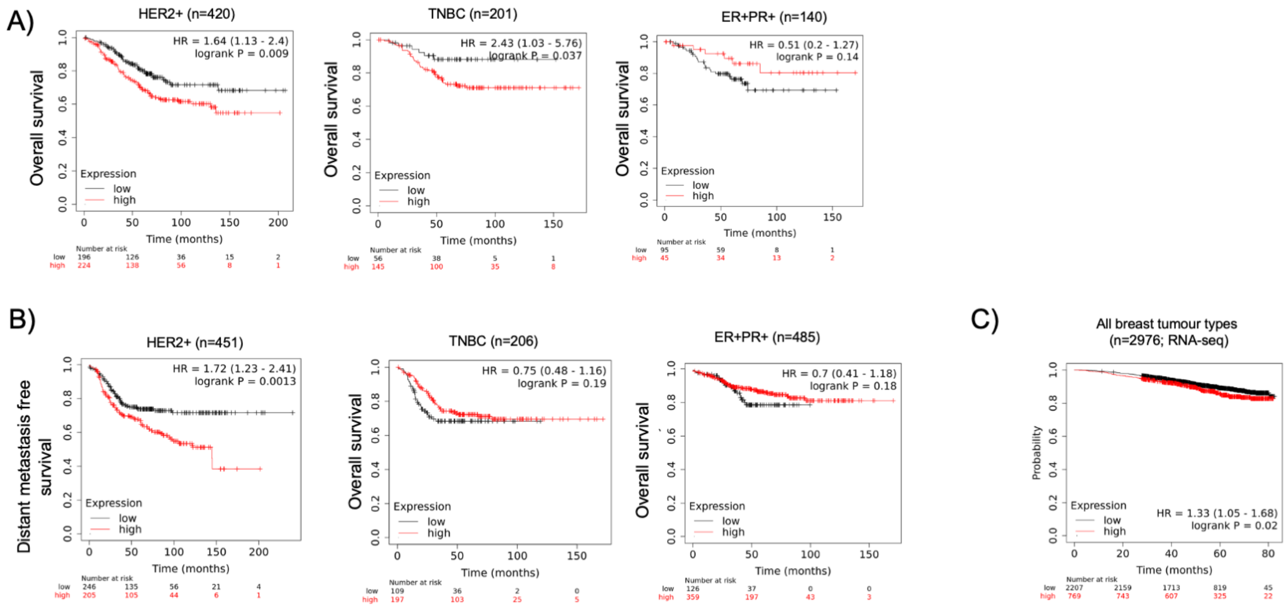

### Supplementary Figure 4

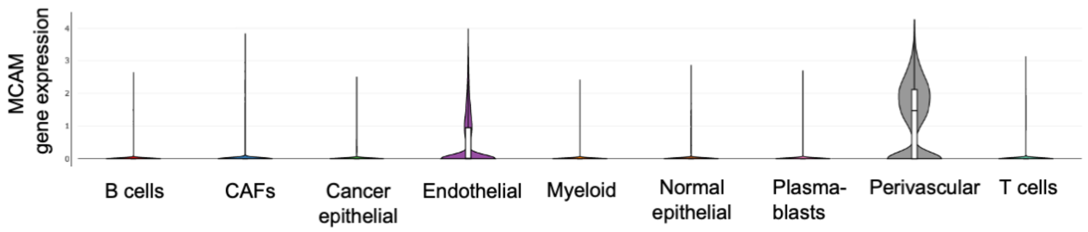


### Supplementary Figure 5

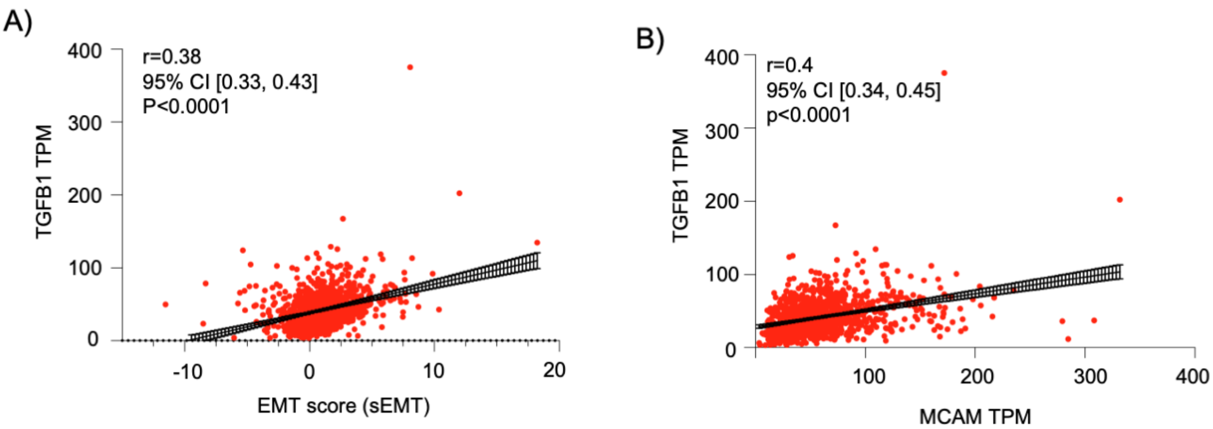


### Supplementary Figure 6

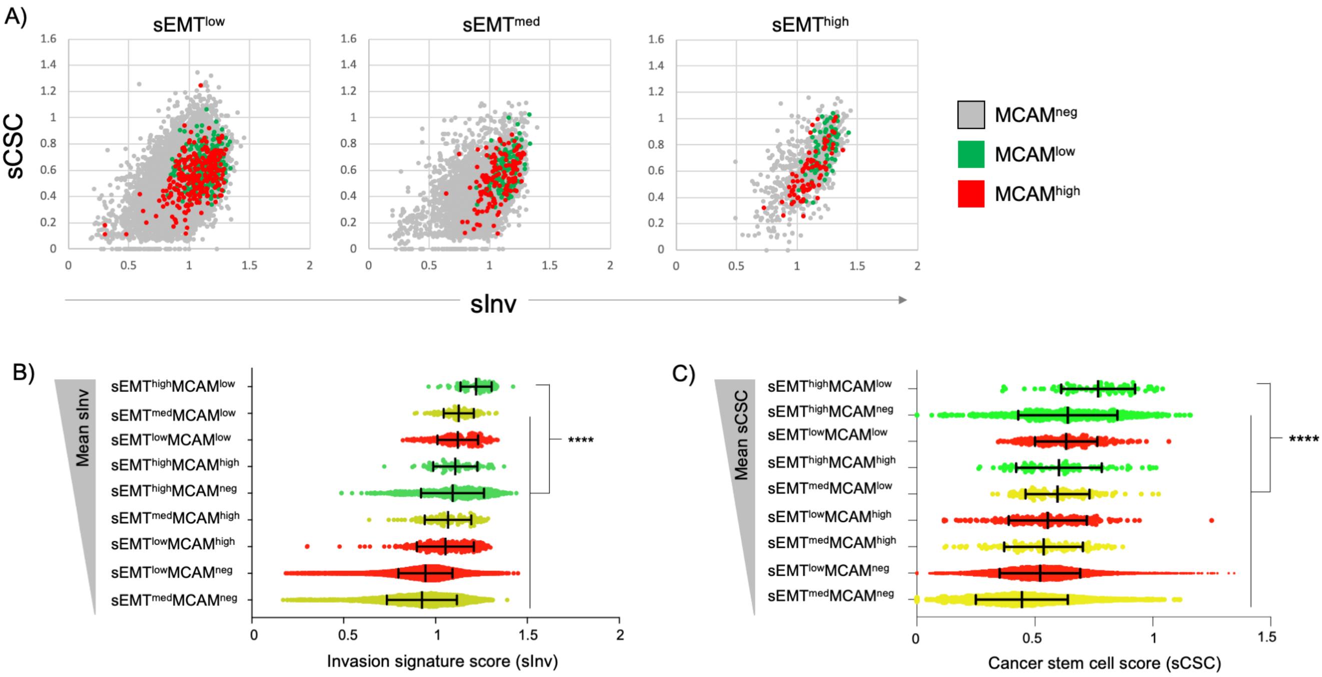

### 2. Supplementary Tables

Supplementary Tables 1 and 2 are attached in a single excel file

**Supplementary Table 1**

CD146/MCAM siRNA sequences

**Supplementary Table 2**

Gene signatures used in this study. The genes used in each signature are listed. The Venn diagram (constructed using https://bioinformatics.psb.ugent.be/webtools/Venn/) indicates that the four signatures are unique, except that MMP1 appears in both sInv and sCSC. The table includes details of how each score was calculated and references.
